# *Ab initio* prediction of specific phospholipid complexes and membrane association of HIV-1 MPER antibodies by multi-scale simulations

**DOI:** 10.1101/2023.05.04.539433

**Authors:** Colleen Maillie, Jay Golden, Ian A. Wilson, Andrew B. Ward, Marco Mravic

**Author notes:** Corresponding author: Marco Mravic.

## Abstract

A potent class of HIV-1 broadly neutralizing antibodies (bnAbs) targets the envelope glycoprotein’s membrane proximal exposed region (MPER) through a proposed mechanism where hypervariable loops embed into lipid bilayers and engage headgroup moieties alongside the epitope. We address the feasibility and determinant molecular features of this mechanism using multi-scale modeling. All-atom simulations of 4E10, PGZL1, 10E8 and LN01 docked onto HIV-like membranes consistently form phospholipid complexes at key complementarity-determining region loop sites, solidifying that stable and specific lipid interactions anchor bnAbs to membrane surfaces. Ancillary protein-lipid contacts reveal surprising contributions from antibody framework regions. Coarse-grained simulations effectively capture antibodies embedding into membranes. Simulations estimating protein-membrane interaction strength for PGZL1 variants along an inferred maturation pathway show bilayer affinity is evolved and correlates with neutralization potency. The modeling demonstrated here uncovers insights into lipid participation in antibodies’ recognition of membrane proteins and highlights antibody features to prioritize in vaccine design.

## Introduction

Antibodies can target integral membrane proteins very near to the lipid bilayer surface, including at epitopes partially embedded within the headgroup region. During antigen engagement, antibodies complementarity-determining regions (CDR) may need to navigate lipid molecules to gain access to a concealed or dynamically buried site. Evolving tolerance or even affinity to lipid bilayers could be beneficial in recognition of membrane-proximal epitopes, potentially forming simultaneous cooperative interactions with antigen and membrane yielding additional avidity and specificity. Conversely, propensity to bind lipids or cell membranes poses a significant auto-immunity risk. B-cells producing antibodies targeting host membranes are downregulated in healthy organisms^1–4^. Nonetheless, rare antibodies with lipid affinity can emerge, particularly in cases of chronic inflammation and infection as in HIV^5–8^. Maturation pathways of these rare events remain unclear, but must strike a careful balance of polyreactivity to avoid or overcome autoreactivity^9^. A better understanding of how antibodies develop membrane affinity and target membrane-proximal epitopes would be impactful for antibody therapeutics, auto-immunity, and vaccine development^10–14^.

We sought to address this phenomenon for broadly neutralizing antibodies (bnAbs) 4E10, PGZL1, 10E8, and LN01 which stem from unique lineages and all target the semi-concealed membrane-proximal epitope region (MPER)^15–18^ of the HIV-1 envelope glycoprotein (Env). Interestingly, these bnAbs show varying degrees of affinity for lipid components; some associate with lipid bilayers or cultured cells even in the absence of antigen^5–819–25^. This membrane interaction behavior appears to correlate with neutralization potency and is attributed to shared CDR loop features including a long hydrophobic CDR-H3^7,20,23,24,26–28^. The correlation between membrane association and neutralization has been established, given that mutations or chemical conjugations to solvent-exposed antibody loop residues which enhance association to phospholipid vesicles consequently boost efficacy of pseudovirus neutralization^29^. Conversely, mutations reducing hydrophobicity drastically reduce neutralization activity while also weakening association to lipid bilayers, despite minimal impacts to antigen affinity (e.g. 4E10 CDR-H3 H100-H102 Trp-Trp motif; PGZL1 clone H4K3’s CDR-H1)^7,23,30,31^. Thus, phospholipid membrane interactions evolved by MPER-targeting bnAbs appear critical for their mechanism of immune protection against HIV. Yet, the breadth of molecular features responsible for association with membranes and mode of lipid-embedded epitope engagement are not clear.

Structural characterization of MPER-targeting bnAbs with full-length Env trimer or gp41 fragments also indicate that the surrounding lipid bilayer plays a role in antibody access and epitope recognition^15,32,33^. Cryo-electron microscopy (cryo-EM) of Env trimers bound to bnAbs PGZL1, 4E10, and 10E8 within in different model membranes suggest their CDR loops form intimate contacts with surrounding lipids while engaging MPER^32,33^. Antigen binding fragments (Fab) crystal structures of those bnAbs as well as LN01 soaked with short-chain phospholipids all revealed ordered headgroup moieties complexed within the CDR loops in the presence and absence of antigen^5,8,22,34,34^ suggesting the antibodies encode specific lipid interactions^5,8,22,34,34^. Thus, the molecular features mediating membrane affinity for these bnAbs appear critical to their maturation and mechanism of immune protection against HIV *in vivo*.

These data support a 2-step bnAb neutralization mechanism proposed previously, wherein a population of bnAbs *in vivo* may first associate with membranes via embedding their CDRs, then laterally diffuse across the bilayer surface to subsequently engage Env at MPER^6,21,24,26,27,32,34–36^. The reverse order, MPER binding followed by CDR membrane insertion, is also often posited. Here, we developed multi-scale molecular dynamics (MD) simulation approaches suited to investigate these mechanisms and the unique maturation landscape these rare antibodies must navigate to avoid auto-immune consequences. For these MPER bnAbs, we focused on the *in-silico* capacity to characterize the molecular features mediating phospholipid affinity at atomic detail, not afforded by previous structural approaches. We find that unbiased all-atom MD simulations accurately and reproducibly predict *ab initio* binding of phospholipids stably bound at specific CDR binding sites previously identified in co-crystal structures^21,22,26,34^. Integrating coarse-grain (CG) simulations, we capture the full process of antibodies scanning and embedding into lipid bilayers to biologically relevant conformations. The simulations illuminate key molecular features tuning bnAbs’ lipid interaction and preferred membrane-bound geometry, advancing on previous work^35^, including surprising contributions from framework regions. Biased simulations provide rough estimates of membrane interaction strength, applied to demonstrate that *in silico* bilayer affinity correlates with neutralization efficacy of experimentally characterized PGZL1 variants along a pseudo-maturation trajectory^18^.

This modeling platform provides an improved framework for navigating the proposed 2-step bnAb neutralization mechanism both for retrospective and proactive evaluation of antibody repertoires, which should have utility in vaccine design and therapeutic antibody engineering. Likewise, our results imply more generalized implications for the *in vivo* selection of antibodies targeting membrane proteins, beyond HIV, at partially concealed juxtamembrane regions – highly desirable epitope regions given their typically high protein sequence conservation. Reliable atomic detail views of antibody conformations at the membrane surface and loop phospholipid binding should be powerful for better understanding membrane protein epitope engagement, pathogen neutralization mechanism, and the checkpoints regulating self-sensing antibodies.

### Atomic simulations accurately model MPER bnAb 4E10 and PGZL1 phospholipid interactions at HIV-like membrane surfaces

To assess the stability, organization, and spectrum of phospholipid interactions inferred from previous lipid-bound crystallography and cryo-EM experiments, we began by performing unbiased all-atom simulations of 4E10, PGZL1, and 10E8 peripherally embedded to the surface of model lipid bilayers. Initial membrane-bound Fab conformations were computed using PPM2.0^3735^, globally optimizing protein insertion based on per-residue hydrophobicity and solvation, each resulting in reasonable predictions with CDR-H3 embedded. Four 1 μs pseudo-replicate simulations were initiated per bnAb, with two replicates tilted by ±15 degrees to modestly vary the starting membrane-interacting pose, utilizing an explicit simplified HIV-like model anionic bilayer: 25% cholesterol, 5% 1-palmitoyl-2-oleoyl-sn-glycero-3-phosphate (POPA), 70% 1-palmitoyl-2-oleoyl-sn-glycero-3-phosphocholine (POPC)^6^.

4E10 and PGZL1 are homologous (85% sequence identity, sharing IgG *VH1-69, VH4-34* and *VH4-59*), and have very similar affinities, breadth, and potency (especially comparing PGZL1.H4K3). 4E10 is notably more poly-reactive: binding to cultured cells, numerous isolated phospholipids^5^, and vesicles of most compositions even in the absence of antigen – whereas 10E8, LN01, and PGZL1 do not. PGZL1 and 4E10 crystal structures both bear putative lipid electron density at several surface loops, notably recurring within a CDR-H1 site – a region when mutation reduced the neutralization potency 10-fold for PGZL1-H4K3^22,34,38^. Given their similarities, we first compared the lipid bilayer interaction mechanisms in 4E10 and PGZL1 simulations. Upon time-averaging phospholipid density from 4 independent trajectories, both bnAbs showed strong hotspots for a lipid phosphate bound within the CDR-H1 loops, with minimal phospholipid or cholesterol ordering around the proteins elsewhere. The simulated lipid phosphates bound within CDR-H1 have exceptional overlap with electron densities and atomic details of modelled headgroups from respective lipid-soaked co-crystal structures of 4E10 and PGZL1^22,26,34^, with phosphate mean positions RMSDs of 0.6 and 0.7 Å, respectively, relative to the X-ray structures (**Figure 1A-B; Figure Supplement 1A-D**). Thus, our simulations reproduce *ab initio* with atomic accuracy the evidence of 4E10 and PGZL1 bnAbs’ mechanism of membrane association. These results validate the CDR-H1 antibody-lipid interactions proposed from previous co-crystal structures are not only feasible in the context of full biologically realistic HIV-like bilayers, but form very readily. The CDR-H1 lipid phosphate complexes appear to be specific interactions that anchor and orient CDR loops at the bilayer surface for antigen engagement, complementing protein-lipid interactions which are less geometrically specific, i.e., lipid tail solvation of apolar groups in the CDR-H loops such as CDR-H3 tryptophans.

**Figure 1.**
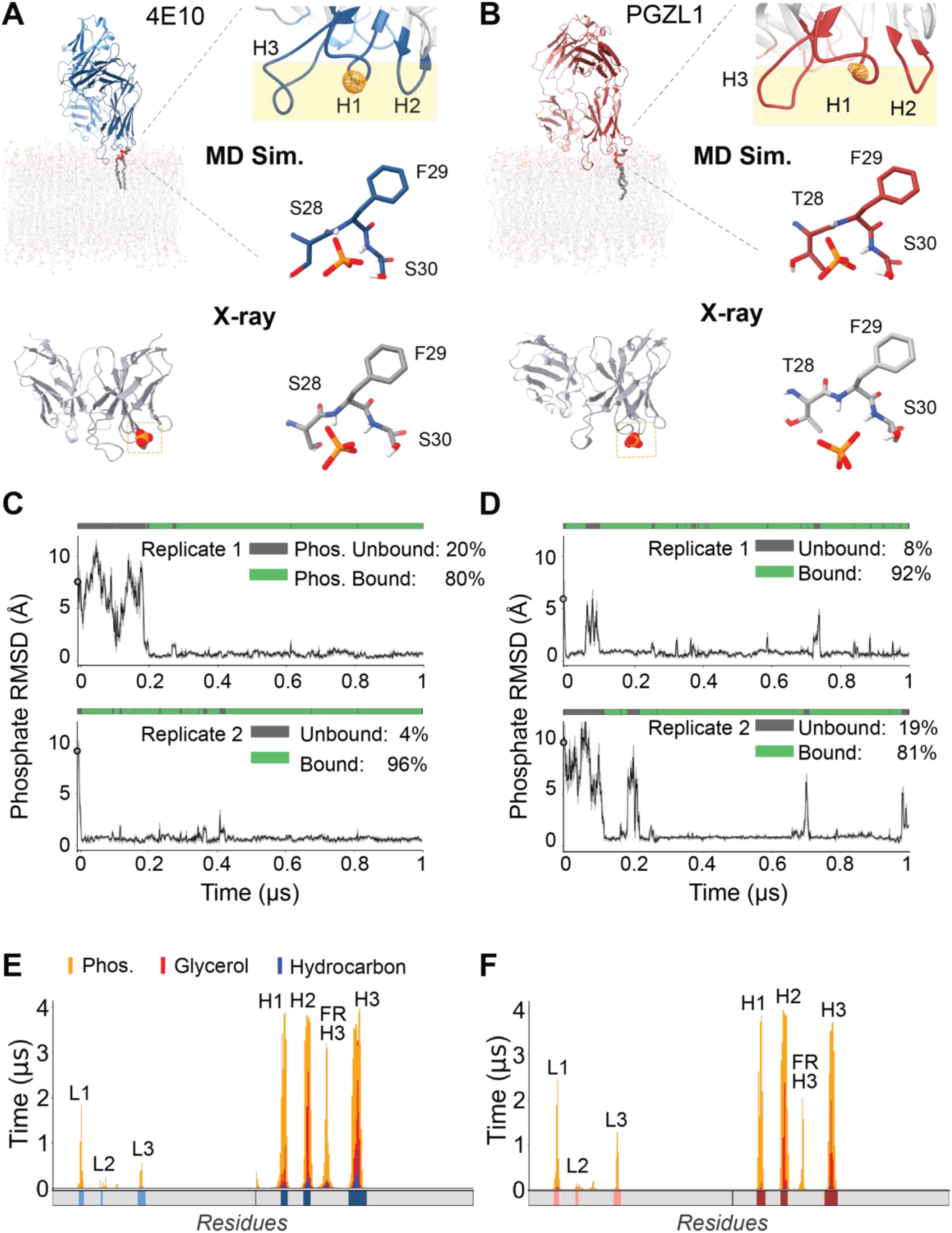
4E10 and PGZL1 CDR lipid phosphate binding sites and membrane interaction profiles in all-atom fluid HIV-like bilayer MD simulations. **(A)** Representative frame from an MD simulation of the phospholipid complex at 4E10 Fab CDR-H1 (top left). Top right, time-averaged lipid phosphate density (orange mesh) relative to antibody CDR loops embedded in the bilayer (beige) from 4 µs total time (top right). Center right, CDR-H1 loop side chain and backbone atoms of *de novo* predicted phosphate interaction (middle right). Bottom, putative lipid phosphate binding site observed in an X-ray structure for 4E10 (PDB: 4XC1), comparing the CDR-H1 loop interactions to MD (bottom right). **(B)** PGZL1 lipid phosphate interaction at the CDR-H1 loop from MD simulation versus X-ray crystallography (PDB: 6O3J), demarked as in (**A**). **(C)** RMSD of the interacting lipid phosphate versus the experimental CDR-phosphate position (X-ray site), classified as bound (green) and unbound (grey) by loop-phosphate contacts (see methods) in 2, 4E10 representative replicate trajectories. Black line, ten-frame RMSD running average; standard deviation, grey shading. **(D)** RMSD of lipid phosphate binding to PGZL1 CDR-H1 in MD simulations versus X-ray site. Phosphate binding is mapped above each MD trajectory as in **(C)**. **(E)** Per-residue interaction profiles for Fab simulations of 4E10 detailing the time spent for each residue in in phosphate layer (orange), glycerol layer (red), or hydrocarbon layer (blue) across aggregate 4 µs from 4 simulations. CDR loops are mapped in solid color blocks below each profile. Fab domain regions making significant contact are labeled, including CDRs and heavy chain framework region 3 (FR-H3). **(F)** Per-residue interaction profiles for antibody Fab simulations for PGZL1, colored as in (**E**).

We next characterized the stability and dynamics of the spontaneously forming antibody-phospholipid complexes which recurred across all trajectories. For both 4E10 and PGZL1, 3 of 4 replicates showed very rapid binding of a headgroup phosphate *ab initio* in the first 20 nanoseconds (ns) from lipids diffusing in from >= 8 Å away) (**Figure 1C-D**). In some cases, phospholipids bound as early as the protein-restrained equilibration (15 ns) where lipids can freely diffuse (**Figure 1—figure supplement 1D-E**). For both 4E10 and PGZL1, one of each replicate simulation had delayed formation of the stable CDR-H1-mediated phospholipid complexes (within 200 ns) requiring minor Fab reorientation on the membrane surface given these replicates’ initialization from an artificially tilted orientation **(Supp. Video 1)**. Upon formation, most phospholipid complexes were highly stable, typically persisting for hundreds of nanoseconds with low mobility of the phosphate headgroup relative to CDR-H1 (<1 Å RMSF) and high overall occupancy (both >80% across 4 microseconds total). An extensive polar network coordinates the lipid phosphate oxygens within the CDR-H1 loop: hydrogen bonds donated from backbone amides of Phe29 and Ser30 and from sidechain hydroxyls of Ser29/Thr29 and Ser30 (**Figure 1C-D**)^22,34^.

Notably, CDR-H loops in contact with the membrane maintained internally rigid backbone conformations during simulation with CDR-H1 having < 1 Å RMSF (**Figure 1–figure supplement 1C-D**). One PGZL1 simulation captured a rare dynamic lipid exchange event (**Figure 1—figure supplement 1E, Supp. Video 2**). A POPC molecule, bound for ∼500 ns, dissociated from the CDR-H1 and was promptly replaced by a second nearby lipid, POPA. These observations, including the kinetic stability and high occupancy, suggest that 4E10 and PGZL1 family antibodies likely are predominantly bound to an anchoring phospholipid when intimately contacting the membrane surface, including during MPER engagement and in the neutralized complex. When CDR-H3 is inserted, CDR-H1 is presented as a rigid pre-formed binding site conveniently at the membrane’s headgroup layer for rapid formation of complexes, with those phospholipids likely in a steady-state equilibrium freely exchanging (nano-microsecond timescale) yet heavily favoring the bound state.

These results of reliably recovering experimentally determined CDR-phospholipid complexes bolsters confidence that antibody behaviors within the simulations are biologically relevant, motivating our deeper analysis of features driving the bnAbs’ protein-lipid interactions and global membrane surface conformations. Characterizing the cumulative per-residue interaction profiles of 4E10 and PGZL1 with bilayer lipids categorized into headgroup, glycerol, or hydrocarbon layers illustrates the deep immersion of heavy-chain loops, with the light-chain making some peripheral polar headgroup contacts (namely CDRs L1 and L3) **(Figure 1E-F)**. The patterns of lipid interactions compare well with previous 4E10 simulations^35,39^, yet are more expansive here – possibly due to the sustained phospholipid-CDR-H1 complex and greater simulation length.

Aligning interaction profiles with corresponding primary sequences contextualizes the amino acid molecular features responsible and their origin in antibody maturation. One novel observation consistent between our 4E10 and PGZL1 simulations was that heavy-chain framework region 3 (FR-H3) extensively intercalates into the bilayer surface. This finding demonstrates that predominantly germline encoded antibody surface features can provide significant contributions to the membrane binding mechanism and surface orientation of antibodies – a key insight for invoking and honing antibody-lipid interactions in vaccine design. For the light-chain loops and FR-H3 of PGZL1 and 4E10, charged residues mediate their shallower headgroup surface interactions **(Figure 1-figure supplement 2A-B)**, whereas the CDR-H loops bare neighboring stretches of hydrophobic and lipophilic polar residues facilitating their deeper embedding. Compared to PGZL1, 4E10’s CDR-H loops are further buried, more extensively contacting lipid aliphatic tails – which may in part explain 4E10’s higher tendency for poly-specificity. The comprehensive protein-lipid atomic-detail maps further detail that the mechanisms for membrane interaction and antibody maturation driven are largely by the solvation for apolar surface loops – the extent of hydrophobic burial – complemented by compatible polar residue electrostatics. The majority of bnAb protein-lipid interactions arise from surface residues’ macroscopic chemical properties, contrasting the much more sequence and conformationally specific phospholipid binding sites constructed at CDR-H1. Thus, the multivalent combination of surface features combined with highly specific lipid anchoring sites may be a convergent theme for maturation and mechanism for MPER-targeting bnAbs, including the previously undervalued contribution from germline-encoded framework regions to lipid association.

### Phospholipid complex formed in the CDR Light Chain groove of 10E8 bound to membrane surfaces

We similarly investigated bnAb 10E8, which differs in its genetic origins and expected light-chain-mediated membrane binding mode^34,36,40,41^. In all 4 replicate simulations, we observed a POPC complexed at a groove between CDR-L1 and FR-L3 **(Figure 2A-B)** which had had significant non-protein electron density in previous 10E8 X-ray structures with and without lipids, which had been modelled as headgroup phosphoglycerol anions, glycerol, or free dihydrogen phosphate^26^. In our simulations, the stably bound POPC phosphate was slightly offset (2.7 Å) from the proposed CDR-L1 crystallographic site **(Figure 2-figure supplement 1B)**, and instead localized to FR-L3 stabilized by a hydrogen bonding network from Ser83 hydroxyl, Gly84 backbone amide, and Asn85 C-alpha proton **(Figure 2A)**. Simultaneously, the POPC choline moiety occupied the precise crystallographic CDR-L1 site, wherein the choline cation was coordinated by four pre-organized unpaired backbone carbonyls stemming from the specific CDR-L1 short loop helix conformation **(Figure 2A)**. We find these bivalent complexes engaging both moieties of the lipid phosphocholine (PC) zwitterion at this CDR-L1 FR-L3 groove with >70% occupancy overall across replicate simulations with high stability: 0.5 Å RMSF for CDR-L1–choline site and 0.5 Å RMSF for FR-L3–phosphate site (**Figure 2-figure supplement 1C-D**). These 10E8 loops were conformationally rigid (RMSF < 1.0 Å, **Figure 2-figure supplement 1E)**, presenting a pre-formed binding site analogous to CDR-H1 in 4E10 and PGZL1. Previous experiments highlight the importance of CDR-L1 phospholipid binding site features in 10E8’s neutralization mechanism, given that the double mutant of R29 and Y38 to alanine exhibited vastly decreased neutralization (>500 fold) across HIV strains with negligible impact to MPER affinity^26^. The time-averaged per-residue lipid interaction profile demonstrates how 10E8 uses both heavy (CDR-H3) and light chain loops (CDRs L1, L2, L3; FR-L3) to orient the antibody geometry on the membrane surface to position 10E8’s phospholipid binding groove site at the headgroup-rich bilayer region **(Figure 2-figure supplement 1F)**. These simulations supplement experimental findings with novel atomic details to establish that genetically distinct MPER antibodies share a consistent mechanism of utilizing CDR loops and framework to mediate bilayer interactions, with 10E8 utilizing FR-L3 for phosphate specificity in a stable lipid complex to anchor this bnAb at the membrane surface.

**Figure 2.**
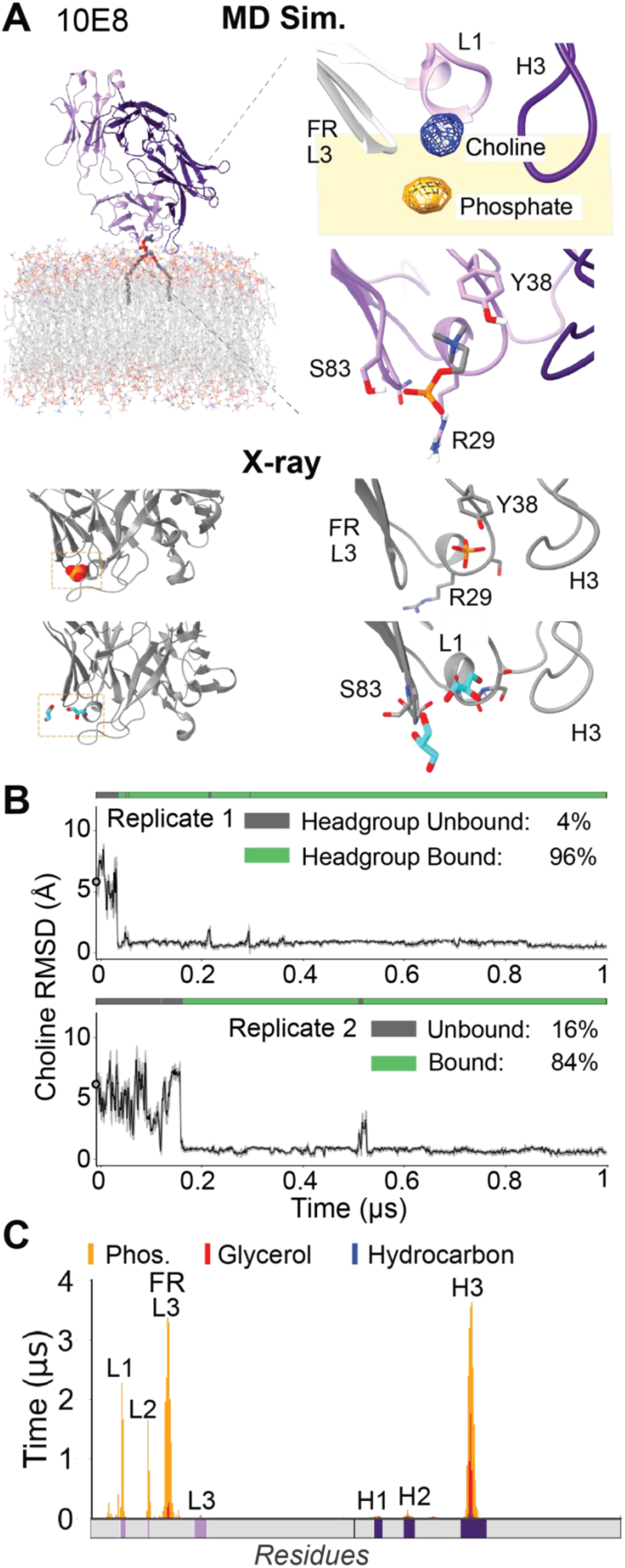
10E8 bivalent lipid headgroup interaction and bilayer insertion predicted in MD simulation closely matches experimental lipid binding sites. **(A)** Top left, representative frame from MD simulation of lipid interacting with 10E8 Fab. Top right, the MD lipid binding site includes a bivalent choline and phosphate lipid headgroup complex (represented as blue and orange mesh time-averaged positional density in simulations) within a protein surface groove composed of CDR-L1 and FR-L3 respectively. Bottom, positions of phosphates or glycerols modeled at the CDR-L1 and FR-L3 groove site within 10E8 Fab X-ray structures (PDB: 5T85, 5T6L) **(B)** RMSD of lipid choline position in the MD simulations versus expected CDR-L1 lipid binding site from 10E8 X-ray structures. “Bound” state assigned relative to choline position and phosphate FR-L3 interactions observed in MD. **(C)** Per-residue interaction profiles for antibody Fab simulations for 10E8 with Fab domain regions making significant contact labeled, including CDRs and light chain framework region 3 (FR-L3).

### Phospholipid binding modes in complex high cholesterol lipid bilayers

We next evaluated whether such bnAb conformations and *ab initio* phospholipid binding are robust to distinct lipid compositions in more complex high cholesterol bilayers matching lipidomics studies for HIV particles^6,42^. New simulations were initialized for similar 4E10, PGZL1, and 10E8 Fab conformations in membranes having a 9.5:19:8.5:18:45 ratio of POPC, 1-palmitoyl-2-oleoyl-sn-glycero-3-phosphoethanolamine (POPE), 1-palmitoyl-2-oleoyl-sn-glycero-3-phospho-L-serine (POPS), palmitoyl sphingomyelin (PSM), Cholesterol. In a 500 ns trajectory for each, stable phospholipid complexes rapidly formed within the respective CDR loop sides with atomic accuracy to the expected position – now having more diversity in lipid headgroup chemistries **(Figure 2–figure supplement 1A-C)**. Although 4E10 bound only POPC at its CDR-H1 site, in PGZL1 simulations CDR-H1 coordinated a PSM molecule (PC headgroup) in precisely the same binding mode as observed for POPC despite PSM’s substitution of two lipid tails off the glycerol backbone. Likewise, 10E8 bound a POPE at its light chain groove site, with the PE (phosphate and cation) forming analogous interactions (<1 Å RMSD to POPC), including coordination of the cationic ethanolamine by the ring of backbone carbonyls capping CDR-L1’s loop helix. Thus, these bnAb phospholipid binding sites can accommodate a broader scope of lipid tail and headgroup chemistries than we have investigated here and can readily form stable phospholipid anchoring complexes in bilayers of HIV-like cholesterol-rich composition. One shared conformational difference observed for these bnAbs the higher cholesterol bilayers was slightly more extensive and broader interaction profiles as well as modestly deeper embedding of the relevant CDR and framework surfaces loops **(Figure 2– figure supplement 1D-F)**. These results bolster the feasibility for using all-atom MD as an in silico platform to explore differential phospholipid affinity at these sites (i.e., specificity studies) and influence on antibody preferred conformation as membrane composition and lipid chemistry are systematically varied. More thorough simulation techniques such as potential of mean force are needed to make those predictions and could better guide experiments concerning how to incorporate the antibody-lipid interaction in neutralization and maturation of bnAbs via vaccine.

### Structural bioinformatics contextualizes bnAb interactions at the membrane

We sought to interrogate the biological relevance and *in silico* predictability of these observed CDR-lipid polar interactions using an orthogonal bioinformatics approach. We mined protein-ligand interactions to assess whether phosphate complexes of similar geometries have been observed in Nature, positing that the occurrence of analogous phosphate interactions outside the context of MPER bnAbs, i.e., within critical structural or functional regions of other protein families, can differentiate common proteome-wide functional motifs from simulation artifacts. Structural searches querying the lipid-binding loop backbone conformation of each Fab (as previously described^16,22,26^) identified between 10^5^ to 2·10^6^ geometrically similar sub-segments within natural proteins (<2 Å RMSD)^43^, reflecting they are relatively prevalent (not rare) in the protein universe, comparing well with frequency of other surface loops of similar length in antibodies (**Supplementary Table 4**). The structurally similar loops found in distinct proteins were subsequently mined for nearby phosphate/phosphoryl and sulfate/sulfo ligands **(Figure 2–figure supplement 3; Supplementary Table 1)**. When searching for loops similar to the CDR-H1 of 4E10 and PGZL1, only 4 cases of phosphate-type ligand binding motifs were identified. Thus, this CDR-H1 site is a realistic but rare protein-phospho-ligand structural motif **(Figure 2–figure supplement 3A-B)**.

For 10E8’s CDR-L1 site, only 1 phosphate-type ligand was observed: a solvent-exposed surface crystallographic ion. By contrast, 10E8’s FR-L3 beta-turn site was a hot spot for phosphate-type ligands, with 23 natural protein cases **(Figure 2–figure supplement 3C)**. Thus, while both CDR-L1 and FR-L3 10E8 loops sites can bind phosphate ligands, the much greater natural prevalence of phosphates ligated at FR-L3 supports that the newly proposed 10E8 POPC complex observed in our simulations is biologically realistic, rather than a simulation artifact. Barring experimental validation, existence of this FR-L3 site establishes precedent for how important framework regions can be for mediating protein-membrane interactions, including hosting stable and specific phospholipid complexes. This bioinformatics analysis illuminates the sophisticated molecular recognition mechanism in which 4E10, PGZL1 and 10E8 Fab domains incorporate recurring structural features from Nature to coordinate specific phospholipid binding interactions.

### Global geometries of membrane-bound bnAb and implications for tilted Env engagement

Given the emphasis on antibody tilt relative to the bilayer and antigen in discerning neutralization mechanisms^32^, we compared the geometries of membrane-bound conformational ensembles for our 4E10, PGZL1, and 10E8 simulations to those of Fab-bound Env trimer cryo-EM structures in membrane mimics^32^. Simulated antigen-free Fabs favor similar orientations on lipid bilayer surfaces as in bnAb-bound tilted Env experimental structures, likely facilitated by the anchoring phospholipid complexes (**Figure 1–figure supplement 1A; Figure 2–figure supplement 1A**). The primary difference is Env-bound Fabs in cryo-EM adopt slightly more shallow approach angles (∼15*°*) relative to the bilayer normal. The simulated bnAbs in isolation prefer slightly more upright orientations, but present CDRs at approximately the same depth and orientation. Thus, these bnAbs appear pre-disposed in their membrane surface conformations, needing only a minor tilt to form the membrane-antibody-antigen neutralization complex. The apparent barrier for re-orientation is likely much less energetically constraining than shielding glycans and accessibility of MPER. Recent studies documenting Env’s inherent stalk flexibility and wide range of ectodomain tilt also suggests looser steric restriction from Env’s soluble domain for an antibody’s bilayer and MPER access than previously thought^33,44^.

Investigating free antibodies’ membrane-bound conformational preferences, e.g., during maturation or for targeting privileged germlines, may be an informative approach to help illicit bnAbs via vaccine with similar lipid-dependent neutralization mechanism. Thus, we characterized the geometric range of 10E8, 4E10, and PGZL1 ensembles within our simulations via structural clustering and analyzed orientation-dependence of their CDR-phospholipid complexes. Fab surface geometries were defined by immersion depths of each CDR loop and 2 global domain angles relative to membrane plane and thresholded to distinguish 7 substates, distributing well into population fractions from 5-35% (**Figure 2–figure supplement 4A-H)**. Clustering of all bnAb simulation time together expectedly resulted in 4E10 and PGZL1 mapping to shared macroscopic states while the more light-chain directed 10E8 clustered separately (**Figure 2–figure supplement 4B-C)**, reflecting the large differences to CDR-H1 depth and approach angle **(Supplementary Table 2).** Upon clustering the 4E10 ensemble, interestingly, all substates maintained 86-98% occupancy of the CDR-H1 phospholipid complex whereas 10E8 and PGZL1 ensemble substates showed more conformation-specific variance in their respective phospholipid binding (**Figure 2–figure supplement 4E,G,I)**. For example, two 10E8 substates exhibit low CDR-L1 PC headgroup occupancy (23%, 33%) due to a domain rotation (20-30*°*) resulting in more CDR-H contacts at the expense of the more typical CDR-L interactions typically observed for 10E8 (**Figure 2–figure supplement 4I**). These findings highlight the important interplay between bnAbs’ preferred surface orientation and loop composition to form these conformationally sensitive anchoring complexes through positioning of the key phospholipid-binding loops. Loop composition and conformational ensembles are likely honed together during the maturation process.

### Antigen influence on membrane bound conformations and lipid binding sites for LN01

We next studied bnAb LN01 to interrogate differences in the antibody surface-bound conformational ensemble and phospholipid interactions in the context of a transmembrane (TM) antigen, utilizing the availability of Fab crystal structures complexed with an MPER-TM fragment **(Figure 3)**^21^. MPER-bound LN01 structures bind one phospholipid and one dodecyl-PC detergent in its CDRs, implying possible paratope-epitope-membrane cooperativity in the neutralization mechanism and two putative phospholipid sites anchoring the bnAb to membrane surfaces (**Figure 3–figure supplement 1A)**. The PC headgroup binding site resides in the CDR-H3-L1-L2 inter-chain groove, with the lipid phosphate engaged by L1 Lys31 via salt bridge and the choline moiety interacting in a cation-pi cage with Tyr32 and CDRH3 Tyr100g. When present, MPER contributes additional choline-aromatic stacking (Trp680, Tyr681). The second more solvent-exposed site lies between CDR-H1 and - H2, ordering PS or PC headgroups in crystal structures. We chose to retain the MPER-TM fragment (668-709) as a monomeric continuous helix from the LN01-bound structure^21^. This conformation has been observed for a similar MPER-TM peptide by NMR^45^ and within bnAb-bound Env by cryo-EM for TM domains in both crossing (dimeric) or separated (monomeric, “tripod”) arrangements^32^. We recognize the caveat that the model MPER-TM antigen fragment studied here does not encompass the full structural breadth of MPER display observed or hypothesized for antigen fragments or full-length Env conformational states. However, alternative MPER-TM organizations observed from peptides (e.g. trimeric TM domain helical bundle, kinked MPER in membrane surface-bound or globular trimer)^46,47^ are less amendable to model due to lack of structural data with bnAbs.

**Figure 3.**
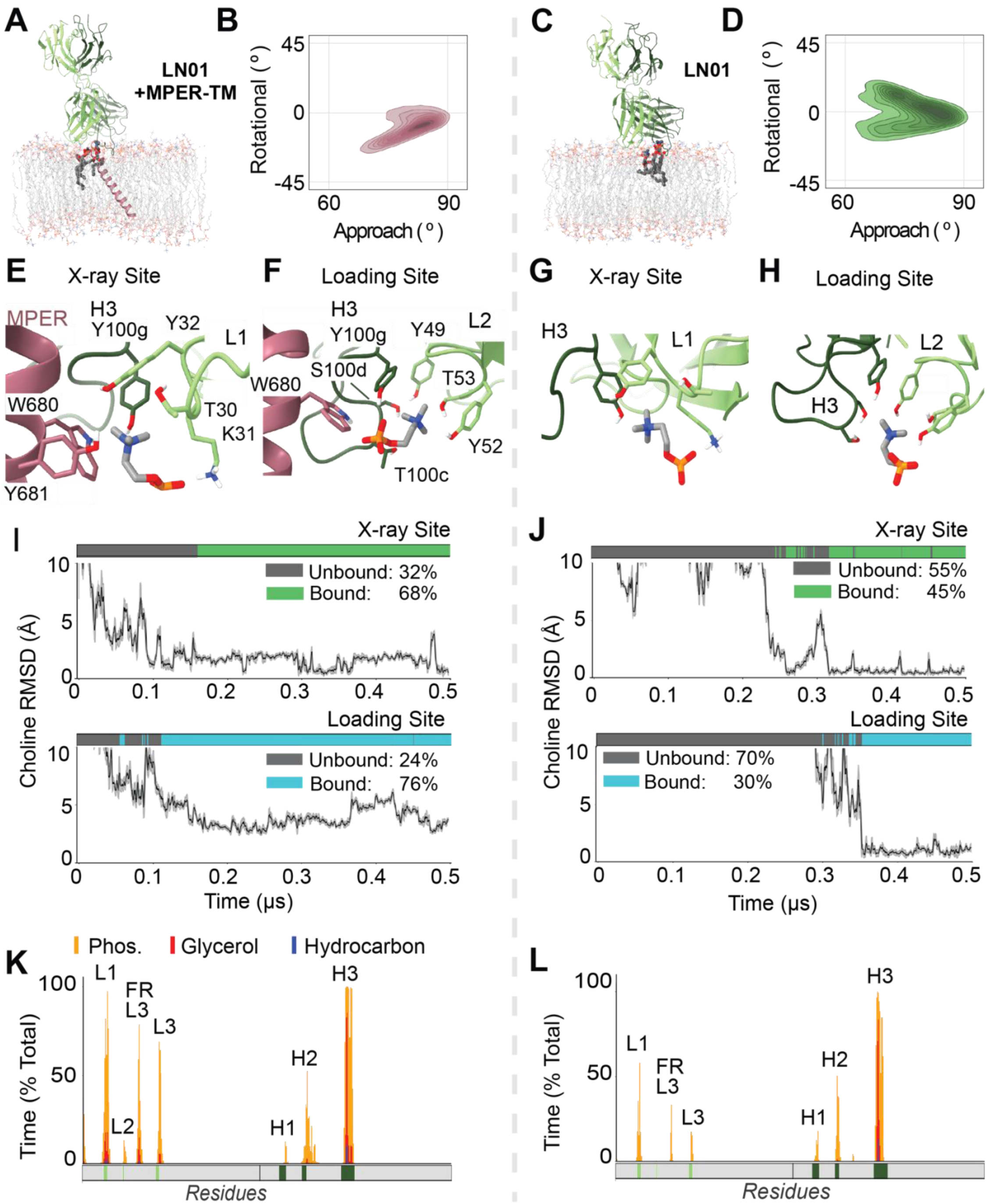
Atomistic simulations of apo and antigen-bound LN01 characterizing paratope-phospholipid complexes with and without epitope and shifted membrane-bound conformation. **(A)** Representative frame from MD simulation LN01 Fab bound to MPER-TM showing the stable ternary paratope-epitope-membrane complex; bound phospholipids shown. **(B)** Frequency of Fab’s characteristic surface-bound geometry by global domain rotation and approach angles in MD simulations for LN01 bound to MPER-TM, plotted by kernel density estimation as contour. **(C)** Representative frame from MD simulation of phospholipids complexed with LN01 Fab alone. **(D)** Frequency of geometries sampled for apo membrane-bound LN01. **(E)** Phospholipid headgroup interaction formed *ab initio* in LN01+MPER-TM simulations. Aromatic cation-pi cage motifs coordinate choline while the phosphate is coordinating by Lys31 matching the X-ray site binding pose. **(F)** The additional distal “Loading” phospholipid site predicted in LN01simulations, with a similar cation-pi cage motif and hydrogen bonds interactions stabilizing the PC headgroup. **(G)** Atomic interactions at the X-ray site in apo LN01 simulations. **(H)** Interactions at the loading site in apo LN01 simulations. **(I)** Lipid headgroup binding in representative simulation of LN01+MPER-TM (n=4 total). Top, X-ray binding site occupancy (green) and phospholipid choline RMSD in a representative trajectory versus experimental position. Bottom, loading site occupancy (cyan) and choline RMSD versus average headgroup bound position. **(J)** Lipid headgroup binding for representative apo LN01 trajectory (n=4 total). Site occupancy and RMSD versus predicted binding position for X-ray site (top) and loading site (bottom). **(K)** Per-residue interaction profile for MPER-TM-bound LN01 aggregated 1.5 µs from 3 simulations. CDR loops are mapped in solid color blocks below each profile. Fab domain regions making significant contact are labeled, including CDRs and light chain framework region 3 (FR-L3). **(L)** Per-residue interaction profile for apo LN01.

Simulations of LN01 were prepared analogously to the other bnAbs with simplified model membranes, replacing 1,2-dioleoyl-sn-glycero-3-phospho-L-serine (DOPS) in place of POPA to provide opportunity to reconstitute LN01’s crystallographic PS binding site. Initial membrane surface geometries for apo and MPER-bound states predicted using PPM^21^, which accounted for solvation of the MPER-TM fragment, lead to similar embedded configurations having CDR-H3 deeply inserted with surface contacts from framework H3, light chain (CDR-L1, L3), and heavy chain (CDR-H1, L2). Across three 1 μs replicates each, those loops remained inserted and adopt ensembles similar to the starting conformations **(Figure 3A-B,K-L)**. The MPER-antibody interface including key interactions (e.g., Trp100H to Thr676 H-bond) remained stable. The TM domain maintains a continuous helix through MPER without kinking in a stable tilted geometry (minimal rotation about helix axis) that enables the previously observed snorkeling and hydration of TM Arg686^48^. In both apo and MPER-TM-bound replicates, several POPC binding events were observed within the CDR-H3-L1-L2 groove at atomic accuracy when compared to dodecyl-PC headgroup position in LN01 X-ray structures, with the same sidechain interactions (**Figure 3E-J)**. In contrast to PGZL1, 4E10, and 10E8 simulations, phospholipids in LN01’s CDR-H3-L1-L2 groove site has much lower kinetic stability and occupancy: ∼30% and ∼40% aggregate occupancy for MPER-TM-bound and apo, respectively (**Figure 3–figure supplement 1C-D**). Occupancy at the second crystallographic CDR-H1-H2 PS/PC site was negligible within any trajectory. Thus, unbiased MD identifies the CDR-H3-L1-L2 groove as the primary phospholipid anchoring site for LN01 in biologically realistic membranes both alone and when bound to TM-embedded MPER, and further emphasizes the importance of lipid in LN01’s maturation and neutralization mechanism.

Two interesting behaviors stood out in LN01 simulations. First, an additional new CDR-phospholipid binding site was observed reproducibly across apo and MPER-TM-bound trajectories at an alternative groove comprised of and CDR-H2, CDR-H3, and FR-L2 we termed the Loading Site (**Figure 3E-J)**. Phospholipids appeared to readily exchange the ∼7 Å distance between this Loading Site and the primary “X-ray” CDR-H3-L1-L2 groove phospholipid binding site (**Figure 3–figure supplement 1B**). Although, the two sites were often simultaneously occupied. In the Loading Site, lipid headgroup phosphate oxygens were coordinated by hydrogen bonds to LN01 Tyr100g, Tyr49, and Tyr52 – and optionally to gp41 Trp680 if present (**Figures 3E-H**). Bivalent interaction with the headgroup choline occurs simultaneously via a similar electrostatic interactions and cation-pi cage motif from Tyr49, Tyr52, Thr53, Thr100c, Ser100d, and Tyr100g sidechains. The Loading site was occupied more often than the X-ray site: 78% and 58% simulation time for MPER-TM-bound and apo LN01 replicates respectively **(Figures 3I,J; Figure 3-figure supplement 1C-D)**.

The second notable behavior observed was that antigen-free LN01 sampled a broader range of geometries at the membrane surface, characterized by two angles describing the Fab relative to the membrane (**Figure 3A-D**). LN01 complexed with MPER-TM was more deeply and extensively embedded than LN01 alone and resulted in a more focused conformation landscape – aligned with the minimal dynamics of the MPER-TM orientation. Protein-lipid interaction profiles reflect consistency in the protein features mediating membrane association between apo and MPER-TM-bound trajectories **(Figure 3K-L)**, with CDR-L and FR-L3 more inserted upon TM antigen engagement upon the minor rearrangement to engage transmembrane antigen. Critically, while apo LN01 favors a different geometry (positive Rotation angle) its membrane surface ensemble heavily samples the major conformation adopted in MPER-TM bound state (negative Rotation angle) (**Figure 3B,D**). This observation that LN01’s surface-bound geometry is predisposed for transmembrane antigen engagement suggests that maturing bnAbs’ sequences and structures are primed for the membrane interaction key to access MPER and form the neuralization complex.

### Coarse-grain *ab initio* insertion simulations capture biologically relevant membrane-bound conformations

Next, we used extended sampling time afforded by Martini coarse-grained (CG) simulations to more thoroughly explore the process of antibody insertion to lipid bilayers and landscape of feasible bnAb surface-bound conformations^49^. Fab’s simulated in CG representation with elastic network restraints maintained the protein fold (<2 Å backbone RMSD) and retained key heavy-light inter-domain contacts, informing that this polar bead model rigidly presents the Fab macroscopic surface features critical for mediating bilayer surface interactions, i.e., hydrophobic patches, charge, polarity (**Figure 5-figure supplement 1A**). Thus, CG simulations might be useful for evaluating bnAbs’ propensity for membrane insertion and approximate preferred conformations, a hypothesis we probed by two distinct simulation approaches.

The first “spontaneous insertion” approach placed a Fab at distinct random initial orientations and distances (0.5 – 2 nm) above a pre-assembled HIV-like anionic lipid bilayer, allowing Fabs free diffusion for 14 µs **(Figure 4A)**. In 18 replicate simulations for each of 4E10, PGZL1, and 10E8, we observed CDR-directed association stable within the lipid bilayer (>1 µs) within 10, 14, and 12 trajectories, respectively (**Figure 4B-D, Supplementary Table 3**). Simulations captured bilayer surface scanning behavior preceding insertion and numerous dissociation events, pointing to the reversibility and dynamics of the process. As a reference, bovine serum albumin (BSA) was tested similarly. BSA’s documented weak affinity for lipid bilayer association (low millimolar)^50^ aligns with the drastically reduced membrane contact events (2/18) and lack of sustained insertion observed **(Figure 4E**). Non-neutralizing anti-gp41 antibody 13h11 having no detectable lipid bilayer interaction by bilayer interferometry^7,51^ also exhibited negligible insertion and only sparse ‘scanning’ events, starkly contrasting the bnAbs’ behaviors. Thus, these CG simulations readily distinguish MPER bnAbs with high propensity for lipid interaction (low micromolar affinity) from non-specific and low affinity lipid interactions.

**Figure 4.**
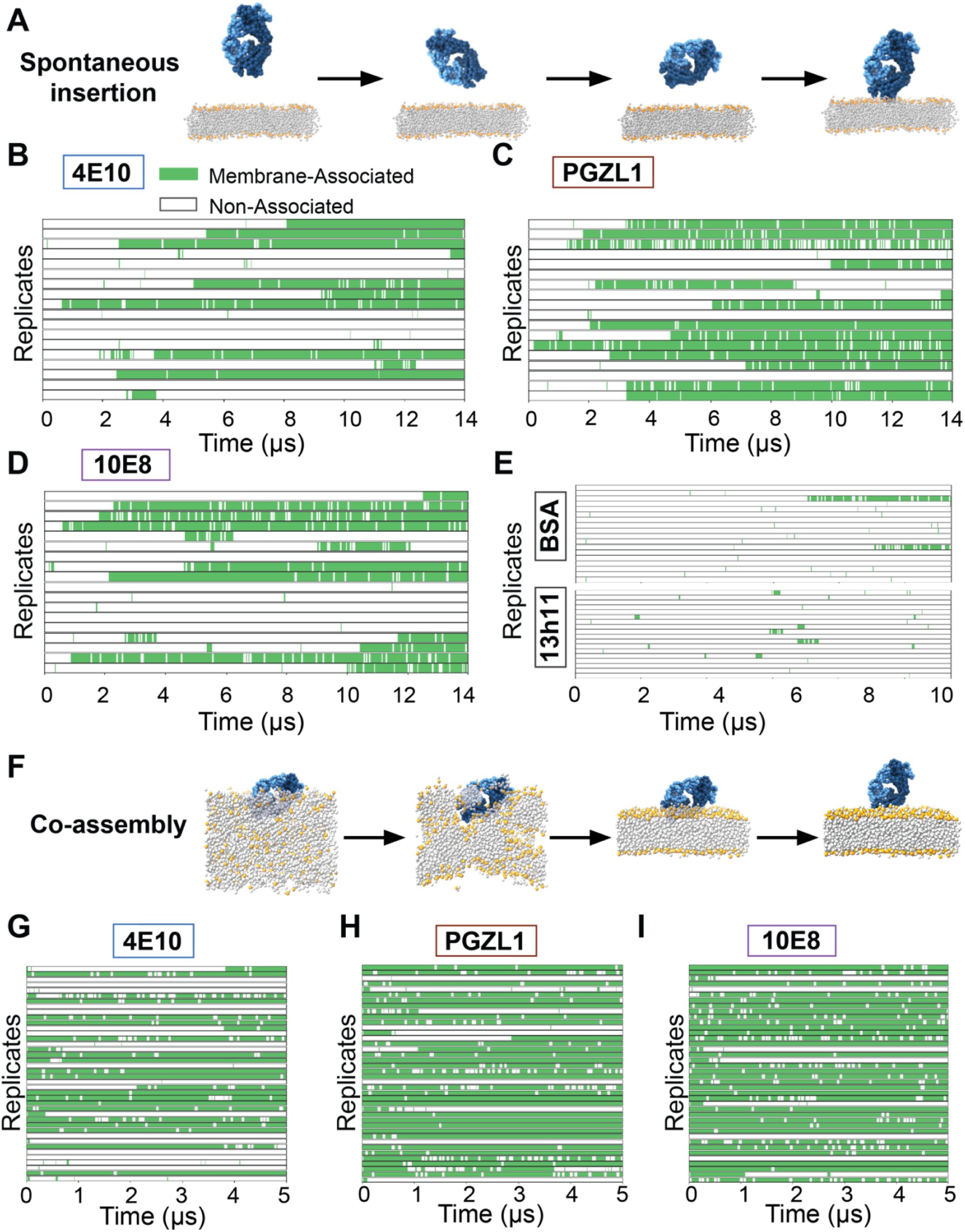
Unbiased spontaneous membrane insertion events and semi-biased dissociation events in coarse grain MD simulations. **(A)** Snapshots of spontaneous insertion event from a Martini model coarse-grain simulation of a 4E10 Fab. The Fab begins in explicit water solvent 0.5-2 nm above a lipid bilayer, freely diffusing and tumbling in bulk solvent, often resulting in a temporary or permanent insertion event (right). **(B,C,D)** 18 replicates of coarse grain Fab systems (4E10, PGZL1, 10E8, respectively), initialized with slightly different Fab orientations relative to lipid bilayer. Frames with Fab contacting the membrane are in green and frames with Fab in water (non-associated) are in white for replicate trajectories of 14 µs each. **(E)** 18 replicates of coarse grain BSA (top) or 13h11 (bottom) with different starting orientations relative to the lipid bilayer. Membrane contact (green) or diffusion in water (white) shown over 10 µs time. **(F)** Snapshots describing a co-assembling membrane pipeline with 4E10 Fab. An Fab is centered in a box with various rotational orientations in space, explicit water, and lipids randomly arranged within a subset of the box (left). By 30ns, the membrane is fully formed (middle). Fab molecules result in a pre-docked membrane bound conformation and sample a permanent insertion event, intermittent membrane association, or dissociation depending on how the Fab contacts with the membrane (right). **(G, H, I)** 40 replicates of 5 µs simulations for coarse grain co-assembling systems (4E10, PGZL1, 10E8, respectively), each with slightly different Fab initial orientations relative to lipid bilayer. Membrane contact is classified as above.

A second “co-assembly” CG approach^52^ aimed to enhance sampling bnAbs insertion and conformations through spontaneous assembly a lipid bilayer around a randomly oriented Fab in a simulation box amongst mixed water, ion, and lipid particles (**Figure 4F**). Predisposition for Fab-lipid interactions and insertion events were increased, particularly for 10E8 and PGZL1, while time wasted sampling Fab in bulk water was reduced. We combated potential biases of this method for possibly enriching insertion in less favorable conformations and skewing distributions by running more replicates (n=40) of shorter 5 µs simulations. Most trajectories having Fabs interacting with lipid with surface features outside CDRs dissociated quickly or never inserted **(Figure 4G-I)**.

We compared the spectrum of membrane-bound conformations sampled from ab initio CG membrane insertion and with pre-inserted all-atom simulations for each bnAb to characterize correlation. Relative Fab orientation was tracked via two global domain angles: the canonical “angle of approach” (Fab long axis intersecting the membrane normal vector) and an internal “rotation angle” from the vector orthogonal to this long axis **(Figure 5A)**. Fab conformational ensembles and protein-lipid interaction patterns from both CG approaches heavily overlapped with the primary geometry observed for each respective bnAb all-atom trajectories, showing consistency between the two simulation models **(Figure 5B, Figure 5-figure supplement 1B-C)**. CG 4E10 insertions across both methods adopted single conformations which were similar and all-atom-like. PGZL1 and 10E8 showed more promiscuity by sampling alternative *ab initio* membrane-inserted geometries alongside their respective all-atom-like conformations, with these states appearing consistently in both CG approaches but at different frequencies. The apparent deeper energy well for 4E10’s primary geometry in CG simulations could reflect greater conformational specificity, but could also be a result of the Martini model’s overestimation of hydrophobic interactions^53^.

**Figure 5.**
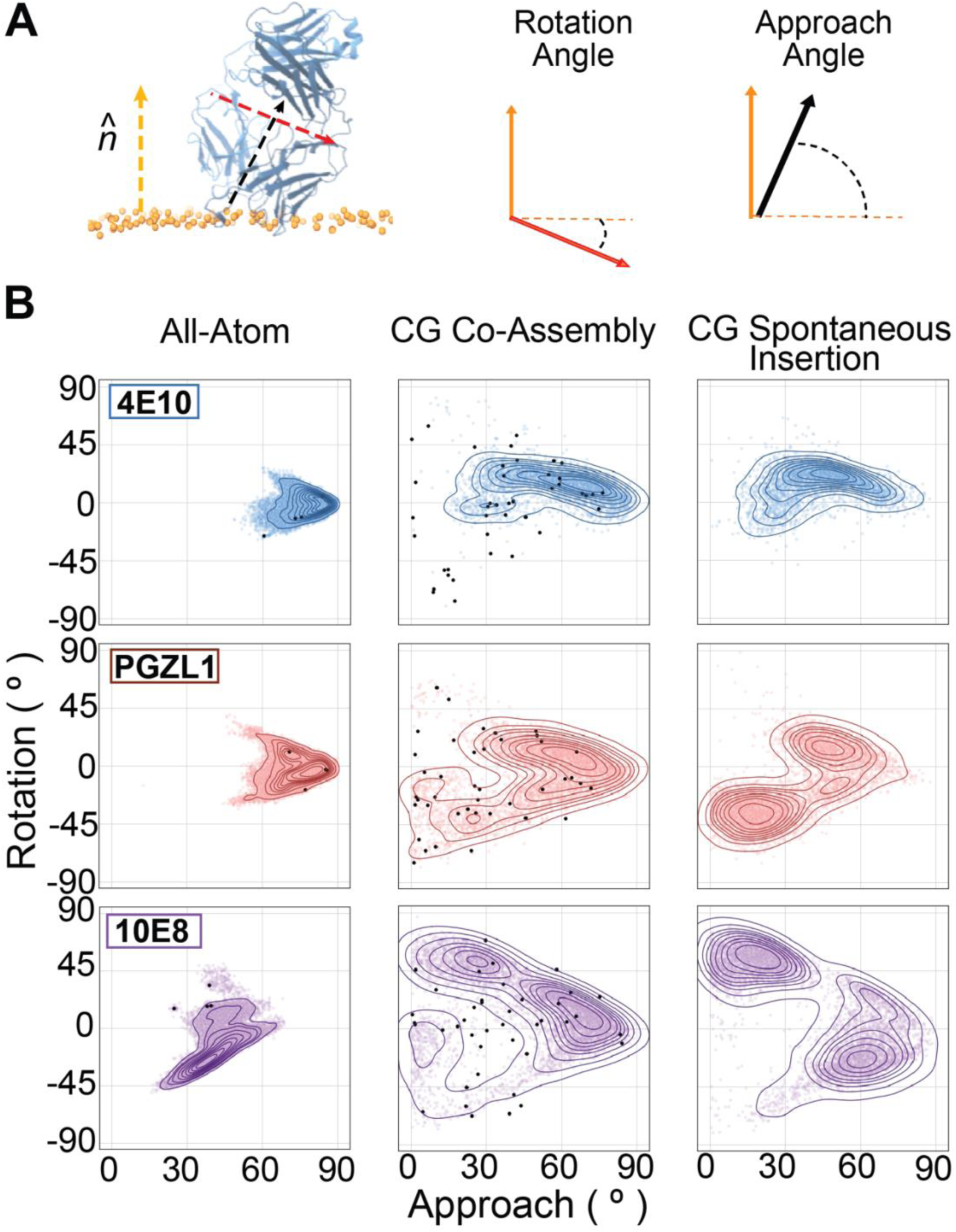
Membrane surface-bound bnAb conformations sampled across multiscale simulations. **(A)** Graphic of angles defined to describe Fab geometries relative to the normal vector at the membrane’s upper leaflet lipid at the phosphate plane in simulation frames (orange arrow). The canonical “approach angle” defines the long axis of the Fab domain (i.e. the central pseudo-symmetry axis) and membrane normal vector (black arrow). A second “rotational angle” is defines the global domain rotation about the Fab pseudo-symmetry axis relative to the membrane normal vector, based on the short axis traversing the light and heavy chains, which is nearly orthogonal to the Fab’s central axis (red arrow). **(B)** Frequency plots of rotation and approach angles from frames of membrane-bound Fabs in MD simulations for 4E10 (blue, top row), PGZL1 (red, middle row), and 10E8 (purple, bottom row). Contour plots depicting frequency maxima for angle pairs sampled are by kernel density estimation. Left column, membrane interaction angles sampled from all-atom simulations with Fabs pre-docked using OPM PPM server prediction. Middle column, geometries from coarse-grain membrane co-assembly simulations. Right column, geometries from unbiased spontaneous insertion coarse-grain simulations. Black dots denote the initial Fab-membrane geometries of starting states for replicate trajectories for each antibody initiated in the lipid bilayer.

Overall, these CG simulations facilitate study of *ab initio* membrane insertion for MPER bnAbs, wherein we find all sustained events are CDR-directed, ruling out contributions from other Fab surface features. For each bnAb, a state we deem to be biologically relevant, based on similarity to geometries observed in all-atom trajectories forming key phospholipid complexes, was either the primary conformation or one of the top 2 sampled. Given that non-CDR directed and alternative CDR-embedded orientations readily dissociate, we conclude that course-grained models can distinguish unfavorable and favorable membrane-bound conformations to an extent that provides utility for characterizing antibody-bilayer interaction mechanisms.

### Multi-scale simulations of ab initio formation of bnAb phospholipid complexes

We next integrated these spontaneously inserted CG conformations into all-atom trajectories to track the full *ab initio* process of bnAb association and determine the competency of CG-derived geometries to subsequently acquire the critical phospholipid anchoring complexes. For each of 4E10, PGZl1, and 10E8, 3 representative medoid Fab poses were extracted by clustering CG simulation frames (clusters A, B, C), and converting to all-atom detail to initiate unbiased MD simulation (**Figure 6A-B**).

**Figure 6.**
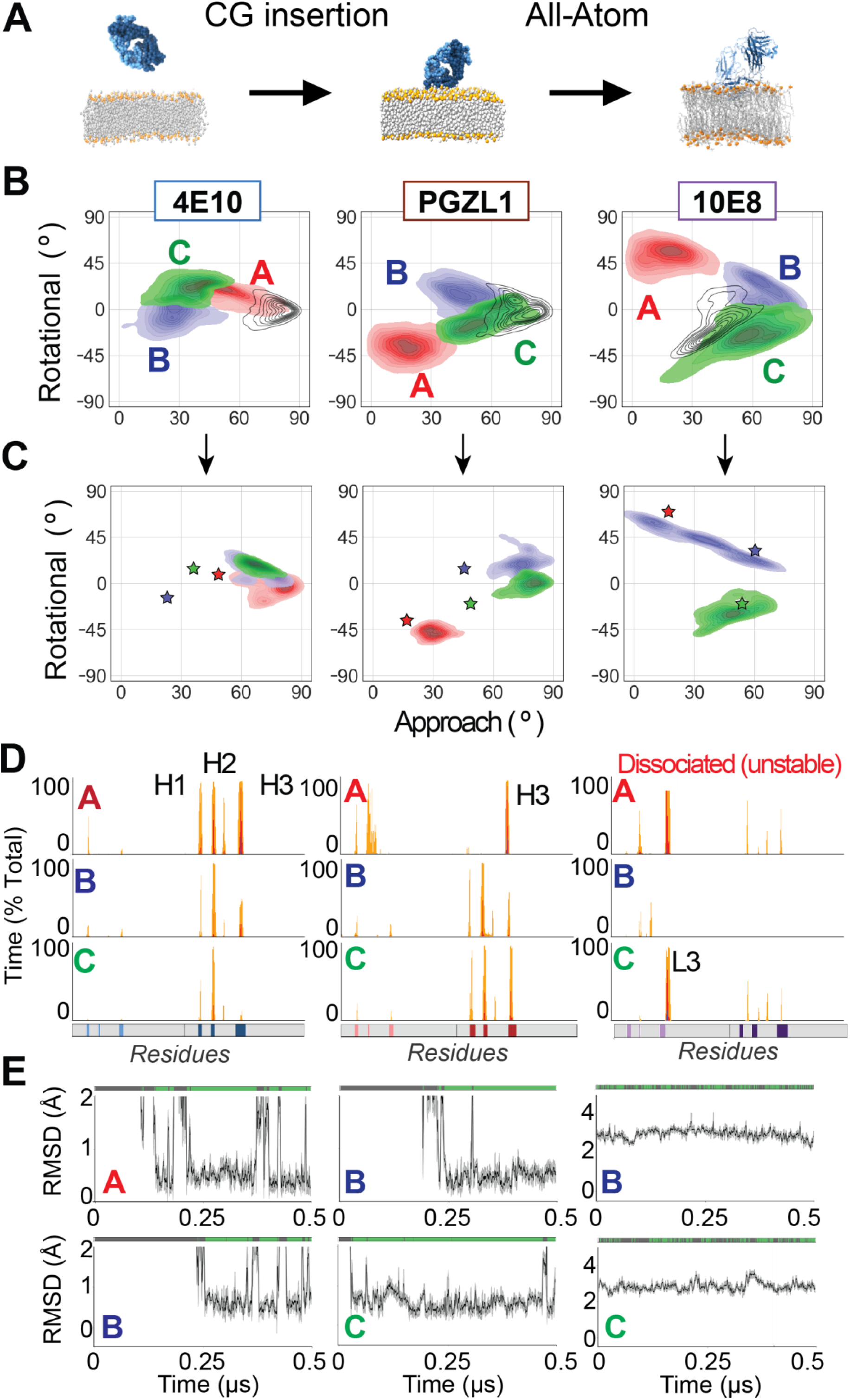
Back-mapping CG membrane-bound geometries to all-atom simulations allows integrative *ab initio* modeling of the full bnAb insertion process. **(A)** Representative frames from membrane-bound 4E10 Fab coarse-grained simulations were back-mapped to all-atom representation to assess the stability and plausibility of those membrane-bound conformations. This coarse-grained-to-all-atom (CG-to-AA) reversion was applied for all medoid frames of interest from each antibody system and used to initiate half-microsecond unbiased all-atom dynamics simulations. **(B)** Frequency of membrane interaction angles from coarse grain spontaneous insertion as clustered by geometric substates for 4E10, PGZL1 and 10E8, colored and contoured as in Fig 3B. Corresponding primary all-atom simulations is overlaid (unfilled, black contour frequency density plot). **(C)** Conformational geometry sampled upon conversion of CG medoid to an all-atom trajectory. Initial geometry denoted by stars colored matching CG clusters in (**B**). Frequency and contour plots of conformational angles sampled in stably inserted backmapped all-atom trajectories for 4E10 (left), PGZL1 (middle), and 10E8 (right). **(D)** Per-residue interaction profiles for antibody Fab simulations for 4E10 (left), PGZL1 (middle), 10E8 (right) representing each backmapped atomic trajectory, showing CDR-mediated conformations of differing depths and geometries. **(E)** Phospholipid headgroup binding and RMSD plots of closest lipid at respective experimentally determined CDR sites for 4E10 (left), PGZL1 (middle), and 10E8 (right), plotted as in Figure 1C or 2B. 4E10 Cluster A is reported in supplement, PGZL1 Cluster A is not reported because no lipid binding was detected, and 10E8 Cluster A is reported in supplement as an artifact.

For 4E10, trajectories initiated from all three geometries drifted back to conformations very similar to those of our initial pre-inserted all-atom trajectories and bound phospholipids at the CDR-H1 site (**Figure 6B-E**). Interestingly, the trajectory back-mapped from cluster A adopted deeper insertion and higher CDR-H1 phospholipid binding occupancy (>60%) compared to ensembles of clusters B and C trajectories, which were slightly tilted (rotation angle > 0°) and less extensively inserted, indicating the latter geometries are likely less ideal (**Figure 6-figure supplement 1B**). For PGZL1, all-atom trajectories starting from CG clusters B and C similarly converge to CDR-H-inserted conformations analogous to our previous all-atom MD, and stably bind phospholipid headgroups at CDR-H1 (>50% occupancy) **(Figure 6C-E)**. For 10E8, Cluster C started and finished in membrane-bound orientations analogous to the pre-inserted simulations (**Figure 6C**); although the Fab adopted a slightly different protein-lipid interaction pattern with much shallower CDR-H3 insertion, the bivalent CDR-L1 phospholipid complex still readily formed with high occupancy (>70%) (**Figure 6D-E).** Thus, these multi-scale simulations demonstrate that unbiased CG modeling can generate stable membrane-bound conformations *ab initio* that integrate smoothly into atomistic simulations which are biologically relevant, in that they capture key phospholipid complexes seen in laboratory experiments inferred to be critical for bnAb function.

CG sampling also explored additional alternative membrane-bound geometries, which we assessed for kinetic stability in integrative atomic simulations to discern secondary protein binding modes or intermediate conformations or aiding membrane association (e.g., preliminary surface-scanning) from simple artifacts. The trajectory initiated from PGZL1’s CG cluster A adopts a novel conformation stable on the 500 ns timescale with CDR-H3 inserted, favoring lipid-CDR-L interactions over CDR-H, that was not competent to complex phospholipids at CDR-H1 (**Figure 6B-D**). All-atom simulation from one alternate conformation of 10E8, CG cluster A, rapidly dissociated from the membrane within 50 ns **(Figure 6-figure supplement 1C)**, showing this integrative MD distinguishes this state as an unstable likely artifact. Interestingly, the all-atom trajectory starting from CG cluster B trended towards the same novel conformation as cluster A, but remained stably inserted and bound phospholipid at the CDR-L1 site at high occupancy (70 %), albeit having shallower CDR-H3 insertion alongside canonical 10E8 CDR-L embedding **(Figure 6C-D)**. These simulations hint at the feasibility to study compatibility of these lipid complex with broader Fab-membrane geometries, and reveal the need for orthogonally evaluating these conformations such as by biased simulations (e.g., potential of mean force, PMF) which we address later in this manuscript.

Additionally, we modelled the orientation of a full-length IgG based on the simulated position of each bnAb Fab to ascertain how sterically feasible a bivalent Fab-membrane interaction could be and possibility of avidity effects in mechanism. The resulting full-length IgG geometries shows that with one membrane-bound Fab, it is very unlikely a second Fabs can simultaneously engage the bilayer due to restriction from the IgG hinge – except for the case an extraordinarily flexible hinge (**Figure 6-figure supplement 1D**).

### Biased pulling simulations distinguish experimentally characterized affinity differences

Finally, we applied biased constant velocity atomistic simulations to compute the force required to dissociate a Fab bound to lipid bilayers – providing a binding strength estimate akin to force spectroscopy **(Figure 7A**). Doing so tested whether this approach could connect the observed bnAb anchoring phospholipid complexes and membrane-bound conformations with physiological properties such as lipid bilayer affinity and neutralization potency. If informative, this method offers an expedient alternative to canonical free energy calculations such as PMFs that demand tens of µs^49^, prohibitive for characterizing many antibody variants. Given that Fabs may sample many lipid interactions and surface conformations, we performed ensemble-based measurements: averaging forces from several replicate pulling trajectories with different starting configurations from unbiased MD.

**Figure 7.**
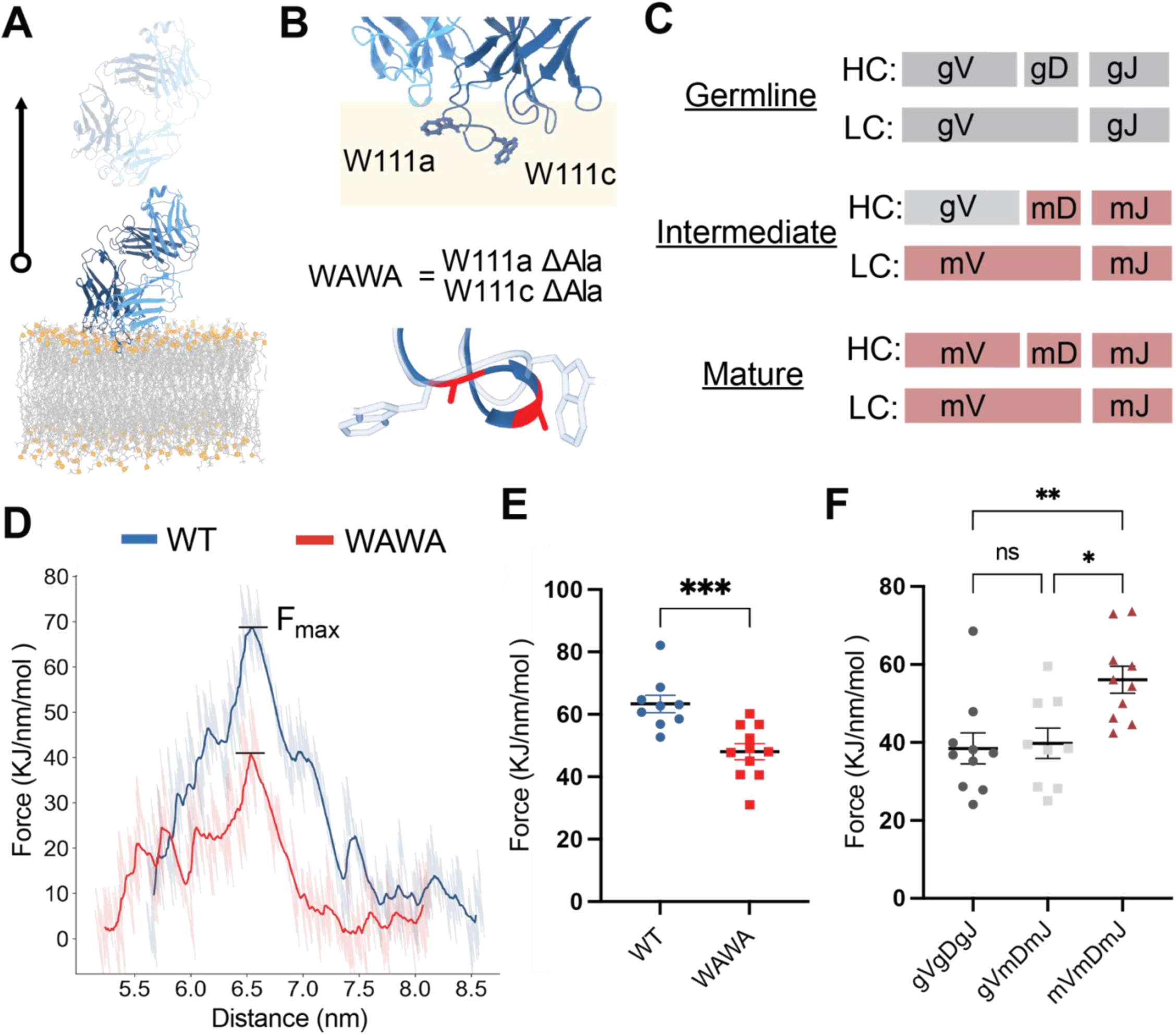
Biased antibody-membrane pulling dissociation simulations approximate Fab-membrane interaction strength for bnAb Fab variants of known lipid binding affinity or neutralization potency. **(A)** Schematic of pulling method, a bnAbs Fabs associated with the lipid bilayers is subject to an applied upward dissociation force to the Fab domain center of mass at constant velocity to measure the force required to dissociate the Fab from the bilayer. **(B)** The Trp100a-Trp100c motif residues in 4E10 CDR-H3 loop of expected to be lipid-embedded in the bilayer (beige). The double alanine mutant “WAWA” 4E10 variant has experimentally determined significantly reduced affinity to lipid bilayers and lower neutralization potency, due to lack of Trp membrane insertion. **(C)** Previously experimentally characterized^20^ PGZL1 germline-reverted variants, shown as chimera of germline versus matured gene segments, used to approximate antibody properties along its maturation trajectory. **(D)** Average force versus distance plots and rupture force (F_max_) calculation for one replicate of pulling wild type 4E10 (WT, blue) and 4E10 WAWA (red) to bias Fab dissociation from the bilayer. **(E)** Distribution of rupture forces required for dissociation for different membrane-bound starting conformations for 4E10 (n=9, blue) and 4E10 WAWA (n=11, red), with starting conformations drawn from previous unbiased all-atom simulation ensembles at rest. Outliers as dots **(F)** Distributions of rupture forces (n=10) required for membrane dissociation of PGZL1 inferred variants along the maturation pathway, for germline (dark grey, gVgDgJ), intermediate (light grey, gVmDmJ), and mature (maroon, mVmDmJ) PGZL1. A one-tailed two-sample t-test was performed to analyze statistical significance of rupture forces between two antibodies (*=p<0.05; **=p<0.005; ***=p<0.001)

We first assessed suitable pulling velocities through dissociating 4E10 poses from simplified anionic HIV-like lipid bilayers to a fixed 1.5 nm distance over different intervals (10, 50, 100, 200 ns; n=3 unique starting conformations). Interestingly, 2 of 3 starting poses adopt the CDR-H3 rotamer conformation of Trp100b projecting down towards lipid tails (“Trp-Down” state) while the third starting pose has Trp100b in a flipped rotamer pointing towards the lipid-water interface (“Trp-Up” state) (**Figure 7-Figure supplement 1A)**. As expected, rupture forces decrease with increasing simulation time (slower pulling velocity), plateauing between 100 and 200 ns (**Figure 7-Figure supplement 1B**). Surprisingly, the rupture forces from trajectories starting in the Trp-Down state were nearly identical (< 2% difference), and rupture forces from trajectories starting from the Trp-Up state were consistently lower (27 and 47% greater) for 50 and 100 ns, respectively. This cursory trial exhibited promising conformational sensitivity in distinguishing different protein-lipid interactions in a CDR loop and precision from similar initial Fab conformations.

We next by compared behavior of 4E10 with its well-studied CDR-H3 WAWA variant, the W100aΔA-W100bΔA double mutant (**Figure 7B**). WAWA has similar K_d_ to MPER as 4E10, but significantly reduced affinity to empty HIV-like liposomes and neutralization efficacy (>100 fold higher IC_50_)^23,27,30,54^. Across replicates from different starting states for 4E10 (n=9) and WAWA (n=11), rupture force calculations showed average forces (± S.E.M.) of 63.8 ± 2.8 kJ/nm/mol and 48.0 ± 2.6 kJ/nm/mol respectively (**Figures 7D-E**), correctly indicating the 4E10 ensemble has a stronger membrane interaction (32 ± 13 %; p-value < 0.001). Thus, this *in silico* estimation effectively discerns antibody variants whose high and low lipid bilayer affinities have been previous measured and reinforces the molecular link between WAWA’s reduced membrane binding and neutralization.

Finally, we evaluated experimentally characterized PGZL1 variants that recapitulate possible stages of a germline maturation pathway^22^ (**Figure 7C**). Mature PGZL1 has relatively high affinity to the MPER epitope peptide (K_d_ = 10 nM) and demonstrates great breadth and potency, neutralizing 84% of a 130-strain panel. An “Intermediate” PGZL1 with V-gene germline reversion (gVmDmJ, 100% CDR-H3 identity) showed a modest affinity loss (K_d_ = 64 nM), while neutralization potency and breadth (12%) were more substantially reduced than expected due to affinity alone – attributed to impaired lipid bilayer interactions^22^. The fully “Germline” reverted variant (gVgDgM) had no detectable MPER peptide affinity or neutralization. Ensembles for Germline and Intermediate Fab pulling simulations were prepared from unbiased all-atom trajectories, starting from predicted hydrophobicity-optimized inserted poses. The variants adopt insertion geometries like the mature Fab, facilitated by the mostly conserved CDR-H3 **(Figure 7-figure supplement 1C)**, but did not form CDR-H1-phospholipid complexes. Significantly lower pulling forces were required to dissociate Intermediate and Germline PGZL1 from bilayers, 37.5 ± 14.9 kJ/nm/mol and 41.2 ± 11.7 kJ/nm/mol respectively, than for mature PGZL1, 56.1 ± 11.6 kJ/nm/mol (p < 0.003) **(Figure 7E)**. Germline PGZL1 likely reflects kinetically stable but low affinity interactions, representing baseline of rupture forces required to dissociate CDR-inserted antibodies. While the difference in Germline and Intermediate force distributions was not significant (p > 0.4), more higher force events were observed for the Intermediate. These results provide further evidence that bnAb phospholipid binding features are acquired and honed along the maturation pathway, entrenching the connection between positive selection of antibody-lipid interactions and neutralization efficacy.

## Discussion

Here, we present a roadmap for applying multi-scale molecular simulations to characterize the biophysical underpinning of lipid membrane interactions within the mechanism of HIV MPER bnAbs. The approaches demonstrated and principles discerned improve understanding of maturation pathways for antibodies targeting membrane-proximal epitopes and should enrich data-driven design of HIV immunogens. Our detailed simulations supplement *in vitro* binding and crystallographic evidence in establishing that bnAbs develop highly specific phospholipid interactions that facilitate access to the MPER epitope and can participate in epitope-paratope interface in context of full lipid bilayers. These results add validation to which out of the many possible ions and short-chain lipid moieties ordered within those crystal structures may be inferred as putative bilayer phospholipids, a critical consideration in antibody and vaccine design. The additional simulations outlined with different lipid ratios and varied antigens as well as more rigorous PMF studies, could be applied to better inform affinity thresholds and energetic relationships paramount to the lipid complexes and conformational preferences reported here. Furthermore, these modeling approaches should have utility in finding membrane-targeting elements in antibodies towards identifying causes of autoimmune stress. Likewise, the analysis tools in this work might be practical for aiding proactive in antibody engineering targeting membrane proteins, for example grafting lipid phosphate binding structural motifs or amino acid sequences with desirable membrane associating properties into distinct antibody loops regions.

Further, these simulations can serve future examination of molecular details concerning the genetic origins and developmental pathways for incipient lipid-binding antibodies. Through natural or vaccine-induced immunity, coaxing the immune system to develop both lipid and antigen affinity (possibly even cooperativity) during maturation is a difficult task, and hindered by downregulation of membrane binding precursors^1,3,6,11^. Likewise, MPER bnAbs are difficult to induce and, until recent clinical trials, have only been isolated from patients with chronic HIV-infection and sustained immunosuppression^13,13,14,19,20,38^. Modern vaccine strategies often elicit certain subsets of precursor B cells, often targeting specific germline genes, in attempt to guide antibodies’ mature of specific molecular features intended for the targeted epitopes^55,56^. The question remains whether particular germline genes and lineages are privileged for successful maturation as membrane-targeting (or MPER-targeting) antibodies. This notion is supported by 4E10 and PGZL1 sharing a germline gene (VH1-69)^22^ and nearly identical membrane interactions. Certain germlines genes may be predisposed with inherent basal membrane affinity or be favored precursor scaffolds for compatibility with evolution of lipid-binding properties. Specific CDR lipid-binding motifs are predicted to be encoded early in maturation processes for PGZL1^22^, namely the CDR-H1 loop, and complemented by mutations at membrane-contacting FR-H3 and CDR-H3 loop residues incorporated later in maturation **(Figure 7-figure supplement 1C)**. Across MPER bnAbs, membrane-binding molecular features are likely acquired at strategic timepoints throughout development to balance poly-specificity and evade autoimmune checkpoints^57^. Experimental databases of germlines genes usage and of B-cell repertoires sequenced during immunization courses would be ideally paired with the simulations described here to investigate *in vivo* filtering of lineages with lipid-binding variants and assess possible rules of the autoimmune system^58^.

Beyond gp41-targeting antibodies and lipid antibodies in autoinflammatory diseases^59,60^ or microbial infections^61,62^, we suspect a broader scope of positive outcomes from antibody tolerance and interaction with lipid bilayers may occur in Nature thus is currently underestimated and underutilized. Although host cross-specificity of 4E10 is well documented, the vastly reduced poly-reactivity of PGZL1 and 10E8 inspires optimism that maturation of phospholipid interactions and membrane tolerance may be incorporated into strategies for vaccine design and therapeutic antibodies for other membrane proteins with conserved buried epitopes^36^. Currently, the breadth of analogous antibodies targeting membrane proteins via extensive or cooperative lipid interactions is poorly explored. The simulation procedures and principles described here are well positioned to investigate this outstanding question and could help define broader chemical rules for design of membrane-interfacing antibodies.

## Acknowledgments

The authors would like to acknowledge the High-Performance Computing Core at Scripps Research and the technical assistance of JC Ducom. Michael B. Zwick and Daniel Leaman provided inspiration and helpful discussions. C.A.M. was supported by the John and Susan Diekman Skaggs Graduate School Fellowship. This work was funded in part by Cooperative Agreement award UM1 AI144462 in partnership with the Division of AIDS, NIAID [I.A.W. and A.B.W.].

## Author contributions

C.A.M. and M.M. performed the MD simulations and conceived of the methodology. C.A.M. quantitatively analyzed the MD simulations. J.G. and C.A.M. conducted the structural bioinformatics analysis. C.A.M., I.A.W., A.B.W., and M.M. designed the experiments, analyzed the data, and wrote the manuscript with input from all authors.

## References

1. Goodnow, C.C., Sprent, J., de St Groth, B.F., and Vinuesa, C.G. (2005). Cellular and genetic mechanisms of self tolerance and autoimmunity. Nature 435, 590–597. 10.1038/nature03724.

2. Liu, M., Yang, G., Wiehe, K., Nicely, N.I., Vandergrift, N.A., Rountree, W., Bonsignori, M., Alam, S.M., Gao, J., Haynes, B.F., et al. (2015). Polyreactivity and Autoreactivity among HIV-1 Antibodies. J Virol 89, 784– 798. 10.1128/JVI.02378-14.

3. Verkoczy, L., Diaz, M., Holl, T.M., Ouyang, Y.-B., Bouton-Verville, H., Alam, S.M., Liao, H.-X., Kelsoe, G., and Haynes, B.F. (2010). Autoreactivity in an HIV-1 broadly reactive neutralizing antibody variable region heavy chain induces immunologic tolerance. Proc. Natl. Acad. Sci. U.S.A. 107, 181–186. 10.1073/pnas.0912914107.

4. Zhang, R., Verkoczy, L., Wiehe, K., Munir Alam, S., Nicely, N.I., Santra, S., Bradley, T., Pemble, C.W., Zhang, J., Gao, F., et al. (2016). Initiation of immune tolerance–controlled HIV gp41 neutralizing B cell lineages. Sci. Transl. Med. 8. 10.1126/scitranslmed.aaf0618.

5. Matyas, G.R., Beck, Z., Karasavvas, N., and Alving, C.R. (2009). Lipid binding properties of 4E10, 2F5, and WR304 monoclonal antibodies that neutralize HIV-1. Biochimica et Biophysica Acta (BBA) - Biomembranes 1788, 660–665. 10.1016/j.bbamem.2008.11.015.

6. Alam, S.M., McAdams, M., Boren, D., Rak, M., Scearce, R.M., Gao, F., Camacho, Z.T., Gewirth, D., Kelsoe, G., Chen, P., et al. (2007). The Role of Antibody Polyspecificity and Lipid Reactivity in Binding of Broadly Neutralizing Anti-HIV-1 Envelope Human Monoclonal Antibodies 2F5 and 4E10 to Glycoprotein 41 Membrane Proximal Envelope Epitopes. The Journal of Immunology 178, 4424–4435. 10.4049/jimmunol.178.7.4424.

7. Alam, S.M., Morelli, M., Dennison, S.M., Liao, H.-X., Zhang, R., Xia, S.-M., Rits-Volloch, S., Sun, L., Harrison, S.C., Haynes, B.F., et al. (2009). Role of HIV membrane in neutralization by two broadly neutralizing antibodies. Proc. Natl. Acad. Sci. U.S.A. 106, 20234–20239. 10.1073/pnas.0908713106.

8. Haynes, B.F., Fleming, J., St. Clair, E.W., Katinger, H., Stiegler, G., Kunert, R., Robinson, J., Scearce, R.M., Plonk, K., Staats, H.F., et al. (2005). Cardiolipin Polyspecific Autoreactivity in Two Broadly Neutralizing HIV-1 Antibodies. Science 308, 1906–1908. 10.1126/science.1111781.

9. Caillat, C., Guilligay, D., Sulbaran, G., and Weissenhorn, W. (2020). Neutralizing Antibodies Targeting HIV-1 gp41. Viruses 12, 1210. 10.3390/v12111210.

10. Burton, D.R., and Hangartner, L. (2016). Broadly Neutralizing Antibodies to HIV and Their Role in Vaccine Design. Annu. Rev. Immunol. 34, 635–659. 10.1146/annurev-immunol-041015-055515.

11. Mascola, J.R., and Haynes, B.F. (2013). HIV-1 neutralizing antibodies: understanding nature’s pathways. Immunol Rev 254, 225–244. 10.1111/imr.12075.

12. Krebs, S.J., Kwon, Y.D., Schramm, C.A., Law, W.H., Donofrio, G., Zhou, K.H., Gift, S., Dussupt, V., Georgiev, I.S., Schätzle, S., et al. (2019). Longitudinal Analysis Reveals Early Development of Three MPER-Directed Neutralizing Antibody Lineages from an HIV-1-Infected Individual. Immunity 50, 677–691.e13. 10.1016/j.immuni.2019.02.008.

13. Rujas, E., Apellániz, B., Torralba, J., Andreu, D., Caaveiro, J.M.M., Wang, S., Lu, S., and Nieva, J.L. (2024). Liposome-based peptide vaccines to elicit immune responses against the membrane active domains of the HIV-1 Env glycoprotein. Biochimica et Biophysica Acta (BBA) - Biomembranes 1866, 184235. 10.1016/j.bbamem.2023.184235.

14. López, C.A., Alam, S.M., Derdeyn, C.A., Haynes, B.F., and Gnanakaran, S. (2024). Influence of membrane on the antigen presentation of the HIV-1 envelope membrane proximal external region (MPER). Current Opinion in Structural Biology 88, 102897. 10.1016/j.sbi.2024.102897.

15. Lee, J.H., Ozorowski, G., and Ward, A.B. (2016). Cryo-EM structure of a native, fully glycosylated, cleaved HIV-1 envelope trimer. Science 351, 1043–1048. 10.1126/science.aad2450.

16. Cardoso, R.M.F., Brunel, F.M., Ferguson, S., Zwick, M., Burton, D.R., Dawson, P.E., and Wilson, I.A. (2007). Structural Basis of Enhanced Binding of Extended and Helically Constrained Peptide Epitopes of the Broadly Neutralizing HIV-1 Antibody 4E10. Journal of Molecular Biology 365, 1533–1544. 10.1016/j.jmb.2006.10.088.

17. Cardoso, R.M.F., Zwick, M.B., Stanfield, R.L., Kunert, R., Binley, J.M., Katinger, H., Burton, D.R., and Wilson, I.A. (2005). Broadly Neutralizing Anti-HIV Antibody 4E10 Recognizes a Helical Conformation of a Highly Conserved Fusion-Associated Motif in gp41. Immunity 22, 163–173. 10.1016/j.immuni.2004.12.011.

18. Sun, Z.-Y.J., Oh, K.J., Kim, M., Yu, J., Brusic, V., Song, L., Qiao, Z., Wang, J., Wagner, G., and Reinherz, E.L. (2008). HIV-1 Broadly Neutralizing Antibody Extracts Its Epitope from a Kinked gp41 Ectodomain Region on the Viral Membrane. Immunity 28, 52–63. 10.1016/j.immuni.2007.11.018.

19. Stiegler, G., Kunert, R., Purtscher, M., Wolbank, S., Voglauer, R., Steindl, F., and Katinger, H. (2001). A Potent Cross-Clade Neutralizing Human Monoclonal Antibody against a Novel Epitope on gp41 of Human Immunodeficiency Virus Type 1. AIDS Research and Human Retroviruses 17, 1757–1765. 10.1089/08892220152741450.

20. Huang, J., Ofek, G., Laub, L., Louder, M.K., Doria-Rose, N.A., Longo, N.S., Imamichi, H., Bailer, R.T., Chakrabarti, B., Sharma, S.K., et al. (2012). Broad and potent neutralization of HIV-1 by a gp41-specific human antibody. Nature 491, 406–412. 10.1038/nature11544.

21. Pinto, D., Fenwick, C., Caillat, C., Silacci, C., Guseva, S., Dehez, F., Chipot, C., Barbieri, S., Minola, A., Jarrossay, D., et al. (2019). Structural Basis for Broad HIV-1 Neutralization by the MPER-Specific Human Broadly Neutralizing Antibody LN01. Cell Host & Microbe 26, 623–637.e8. 10.1016/j.chom.2019.09.016.

22. Zhang, L., Irimia, A., He, L., Landais, E., Rantalainen, K., Leaman, D.P., Vollbrecht, T., Stano, A., Sands, D.I., Kim, A.S., et al. (2019). An MPER antibody neutralizes HIV-1 using germline features shared among donors. Nat Commun 10, 5389. 10.1038/s41467-019-12973-1.

23. Scherer, E.M., Leaman, D.P., Zwick, M.B., McMichael, A.J., and Burton, D.R. (2010). Aromatic residues at the edge of the antibody combining site facilitate viral glycoprotein recognition through membrane interactions. Proc. Natl. Acad. Sci. U.S.A. 107, 1529–1534. 10.1073/pnas.0909680107.

24. Chen, J., Frey, G., Peng, H., Rits-Volloch, S., Garrity, J., Seaman, M.S., and Chen, B. (2014). Mechanism of HIV-1 Neutralization by Antibodies Targeting a Membrane-Proximal Region of gp41. J Virol 88, 1249–1258. 10.1128/JVI.02664-13.

25. Yang, G., Holl, T.M., Liu, Y., Li, Y., Lu, X., Nicely, N.I., Kepler, T.B., Alam, S.M., Liao, H.-X., Cain, D.W., et al. (2013). Identification of autoantigens recognized by the 2F5 and 4E10 broadly neutralizing HIV-1 antibodies. Journal of Experimental Medicine 210, 241–256. 10.1084/jem.20121977.

26. Irimia, A., Serra, A.M., Sarkar, A., Jacak, R., Kalyuzhniy, O., Sok, D., Saye-Francisco, K.L., Schiffner, T., Tingle, R., Kubitz, M., et al. (2017). Lipid interactions and angle of approach to the HIV-1 viral membrane of broadly neutralizing antibody 10E8: Insights for vaccine and therapeutic design. PLoS Pathog 13, e1006212. 10.1371/journal.ppat.1006212.

27. Kwon, Y.D., Chuang, G.-Y., Zhang, B., Bailer, R.T., Doria-Rose, N.A., Gindin, T.S., Lin, B., Louder, M.K., McKee, K., O’Dell, S., et al. (2018). Surface-Matrix Screening Identifies Semi-Specific Interactions that Improve Potency of a Near Pan-reactive HIV-1-Neutralizing Antibody. Cell Reports 22, 1798–1809. 10.1016/j.celrep.2018.01.023.

28. Zwick, M.B., Komori, H.K., Stanfield, R.L., Church, S., Wang, M., Parren, P.W.H.I., Kunert, R., Katinger, H., Wilson, I.A., and Burton, D.R. (2004). The Long Third Complementarity-Determining Region of the Heavy Chain Is Important in the Activity of the Broadly Neutralizing Anti-Human Immunodeficiency Virus Type 1 Antibody 2F5. J Virol 78, 3155–3161. 10.1128/JVI.78.6.3155-3161.2004.

29. Rujas, E., Leaman, D.P., Insausti, S., Ortigosa-Pascual, L., Zhang, L., Zwick, M.B., and Nieva, J.L. (2018). Functional Optimization of Broadly Neutralizing HIV-1 Antibody 10E8 by Promotion of Membrane Interactions. J Virol 92, e02249–17. 10.1128/JVI.02249-17.

30. Xu, H., Song, L., Kim, M., Holmes, M.A., Kraft, Z., Sellhorn, G., Reinherz, E.L., Stamatatos, L., and Strong, R.K. (2010). Interactions between Lipids and Human Anti-HIV Antibody 4E10 Can Be Reduced without Ablating Neutralizing Activity. J Virol 84, 1076–1088. 10.1128/JVI.02113-09.

31. Julien, J.-P., Huarte, N., Maeso, R., Taneva, S.G., Cunningham, A., Nieva, J.L., and Pai, E.F. (2010). Ablation of the Complementarity-Determining Region H3 Apex of the Anti-HIV-1 Broadly Neutralizing Antibody 2F5 Abrogates Neutralizing Capacity without Affecting Core Epitope Binding. J Virol 84, 4136–4147. 10.1128/JVI.02357-09.

32. Rantalainen, K., Berndsen, Z.T., Antanasijevic, A., Schiffner, T., Zhang, X., Lee, W.-H., Torres, J.L., Zhang, L., Irimia, A., Copps, J., et al. (2020). HIV-1 Envelope and MPER Antibody Structures in Lipid Assemblies. Cell Reports 31, 107583. 10.1016/j.celrep.2020.107583.

33. Yang, S., Hiotis, G., Wang, Y., Chen, J., Wang, J., Kim, M., Reinherz, E.L., and Walz, T. (2022). Dynamic HIV-1 spike motion creates vulnerability for its membrane-bound tripod to antibody attack. Nat Commun 13, 6393. 10.1038/s41467-022-34008-y.

34. Irimia, A., Sarkar, A., Stanfield, R.L., and Wilson, I.A. (2016). Crystallographic Identification of Lipid as an Integral Component of the Epitope of HIV Broadly Neutralizing Antibody 4E10. Immunity 44, 21–31. 10.1016/j.immuni.2015.12.001.

35. Carravilla, P., Darré, L., Oar-Arteta, I.R., Vesga, A.G., Rujas, E., de las Heras-Martínez, G., Domene, C., Nieva, J.L., and Requejo-Isidro, J. (2020). The Bilayer Collective Properties Govern the Interaction of an HIV-1 Antibody with the Viral Membrane. Biophysical Journal 118, 44–56. 10.1016/j.bpj.2019.11.005.

36. Soto, C., Ofek, G., Joyce, M.G., Zhang, B., McKee, K., Longo, N.S., Yang, Y., Huang, J., Parks, R., Eudailey, J., et al. (2016). Developmental Pathway of the MPER-Directed HIV-1-Neutralizing Antibody 10E8. PLoS One 11, e0157409. 10.1371/journal.pone.0157409.

37. Lomize, M.A., Pogozheva, I.D., Joo, H., Mosberg, H.I., and Lomize, A.L. (2012). OPM database and PPM web server: resources for positioning of proteins in membranes. Nucleic Acids Research 40, D370–D376. 10.1093/nar/gkr703.

38. Zwick, M.B., Labrijn, A.F., Wang, M., Spenlehauer, C., Saphire, E.O., Binley, J.M., Moore, J.P., Stiegler, G., Katinger, H., Burton, D.R., et al. (2001). Broadly Neutralizing Antibodies Targeted to the Membrane-Proximal External Region of Human Immunodeficiency Virus Type 1 Glycoprotein gp41. J Virol 75, 10892– 10905. 10.1128/JVI.75.22.10892-10905.2001.

39. Carravilla, P., Nieva, J.L., and Eggeling, C. (2020). Fluorescence Microscopy of the HIV-1 Envelope. Viruses 12, 348. 10.3390/v12030348.

40. Georgiev, I.S., Rudicell, R.S., Saunders, K.O., Shi, W., Kirys, T., McKee, K., O’Dell, S., Chuang, G.-Y., Yang, Z.-Y., Ofek, G., et al. (2014). Antibodies VRC01 and 10E8 Neutralize HIV-1 with High Breadth and Potency Even with Ig-Framework Regions Substantially Reverted to Germline. The Journal of Immunology 192, 1100–1106. 10.4049/jimmunol.1302515.

41. Williams, L.D., Ofek, G., Schätzle, S., McDaniel, J.R., Lu, X., Nicely, N.I., Wu, L., Lougheed, C.S., Bradley, T., Louder, M.K., et al. (2017). Potent and broad HIV-neutralizing antibodies in memory B cells and plasma. Sci. Immunol. 2, eaal2200. 10.1126/sciimmunol.aal2200.

42. Brügger, B., Glass, B., Haberkant, P., Leibrecht, I., Wieland, F.T., and Kräusslich, H.-G. (2006). The HIV lipidome: A raft with an unusual composition. Proc. Natl. Acad. Sci. U.S.A. 103, 2641–2646. 10.1073/pnas.0511136103.

43. Polizzi, N.F., and DeGrado, W.F. (2020). A defined structural unit enables de novo design of small-molecule–binding proteins. Science 369, 1227–1233. 10.1126/science.abb8330.

44. Mangala Prasad, V., Leaman, D.P., Lovendahl, K.N., Croft, J.T., Benhaim, M.A., Hodge, E.A., Zwick, M.B., and Lee, K.K. (2022). Cryo-ET of Env on intact HIV virions reveals structural variation and positioning on the Gag lattice. Cell 185, 641–653.e17. 10.1016/j.cell.2022.01.013.

45. Chiliveri, S.C., Louis, J.M., Ghirlando, R., Baber, J.L., and Bax, A. (2018). Tilted, Uninterrupted, Monomeric HIV-1 gp41 Transmembrane Helix from Residual Dipolar Couplings. J. Am. Chem. Soc. 140, 34–37. 10.1021/jacs.7b10245.

46. Piai, A., Fu, Q., Sharp, A.K., Bighi, B., Brown, A.M., and Chou, J.J. (2021). NMR Model of the Entire Membrane-Interacting Region of the HIV-1 Fusion Protein and Its Perturbation of Membrane Morphology. J. Am. Chem. Soc. 143, 6609–6615. 10.1021/jacs.1c01762.

47. Kwon, B., Lee, M., Waring, A.J., and Hong, M. (2018). Oligomeric Structure and Three-Dimensional Fold of the HIV gp41 Membrane-Proximal External Region and Transmembrane Domain in Phospholipid Bilayers. J. Am. Chem. Soc. 140, 8246–8259. 10.1021/jacs.8b04010.

48. Hollingsworth, L.R., Lemkul, J.A., Bevan, D.R., and Brown, A.M. (2018). HIV-1 Env gp41 Transmembrane Domain Dynamics Are Modulated by Lipid, Water, and Ion Interactions. Biophysical Journal 115, 84–94. 10.1016/j.bpj.2018.05.022.

49. Corey, R.A., Vickery, O.N., Sansom, M.S.P., and Stansfeld, P.J. (2019). Insights into Membrane Protein–Lipid Interactions from Free Energy Calculations. J. Chem. Theory Comput. 15, 5727–5736. 10.1021/acs.jctc.9b00548.

50. Ruggeri, F., Zhang, F., Lind, T., Bruce, E.D., Lau, B.L.T., and Cárdenas, M. (2013). Non-specific interactions between soluble proteins and lipids induce irreversible changes in the properties of lipid bilayers. Soft Matter 9, 4219–4226. 10.1039/C3SM27769K.

51. Nicely, N.I., Dennison, S.M., Spicer, L., Scearce, R.M., Kelsoe, G., Ueda, Y., Chen, H., Liao, H.-X., Alam, S.M., and Haynes, B.F. (2010). Crystal structure of a non-neutralizing antibody to the HIV-1 gp41 membrane-proximal external region. Nat Struct Mol Biol 17, 1492–1494. 10.1038/nsmb.1944.

52. Scott, K.A., Bond, P.J., Ivetac, A., Chetwynd, A.P., Khalid, S., and Sansom, M.S.P. (2008). Coarse-Grained MD Simulations of Membrane Protein-Bilayer Self-Assembly. Structure 16, 621–630. 10.1016/j.str.2008.01.014.

53. Srinivasan, S., Zoni, V., and Vanni, S. (2021). Estimating the accuracy of the MARTINI model towards the investigation of peripheral protein–membrane interactions. Faraday Discuss. 232, 131–148. 10.1039/D0FD00058B.

54. Montero, M., Gulzar, N., Klaric, K.-A., Donald, J.E., Lepik, C., Wu, S., Tsai, S., Julien, J.-P., Hessell, A.J., Wang, S., et al. (2012). Neutralizing Epitopes in the Membrane-Proximal External Region of HIV-1 gp41 Are Influenced by the Transmembrane Domain and the Plasma Membrane. J Virol 86, 2930–2941. 10.1128/JVI.06349-11.

55. Jardine, J., Julien, J.-P., Menis, S., Ota, T., Kalyuzhniy, O., McGuire, A., Sok, D., Huang, P.-S., MacPherson, S., Jones, M., et al. (2013). Rational HIV Immunogen Design to Target Specific Germline B Cell Receptors. Science 340, 711–716. 10.1126/science.1234150.

56. Steichen, J.M., Kulp, D.W., Tokatlian, T., Escolano, A., Dosenovic, P., Stanfield, R.L., McCoy, L.E., Ozorowski, G., Hu, X., Kalyuzhniy, O., et al. (2016). HIV Vaccine Design to Target Germline Precursors of Glycan-Dependent Broadly Neutralizing Antibodies. Immunity 45, 483–496. 10.1016/j.immuni.2016.08.016.

57. Klein, F., Diskin, R., Scheid, J.F., Gaebler, C., Mouquet, H., Georgiev, I.S., Pancera, M., Zhou, T., Incesu, R.-B., Fu, B.Z., et al. (2013). Somatic Mutations of the Immunoglobulin Framework Are Generally Required for Broad and Potent HIV-1 Neutralization. Cell 153, 126–138. 10.1016/j.cell.2013.03.018.

58. Briney, B., Inderbitzin, A., Joyce, C., and Burton, D.R. (2019). Commonality despite exceptional diversity in the baseline human antibody repertoire. Nature 566, 393–397. 10.1038/s41586-019-0879-y.

59. Dema, B., and Charles, N. (2016). Autoantibodies in SLE: Specificities, Isotypes and Receptors. Antibodies 5, 2. 10.3390/antib5010002.

60. Tao, T.-W., and Kriss, J.P. (1982). Membrane-Binding Antibodies in Patients with Graves’ Disease and Other Autoimmune Diseases*. The Journal of Clinical Endocrinology & Metabolism 55, 935–940. 10.1210/jcem-55-5-935.

61. Asherson, R.A. (2003). Antiphospholipid antibodies and infections. Annals of the Rheumatic Diseases 62, 388–393. 10.1136/ard.62.5.388.

62. Sène, D., Piette, J.-C., and Cacoub, P. (2008). Antiphospholipid antibodies, antiphospholipid syndrome and infections. Autoimmunity Reviews 7, 272–277. 10.1016/j.autrev.2007.10.001.

