## Supplemental Figures for "*Ab initio* prediction of specific phospholipid complexes and membrane association of HIV-1 MPER antibodies by multi-scale simulations"

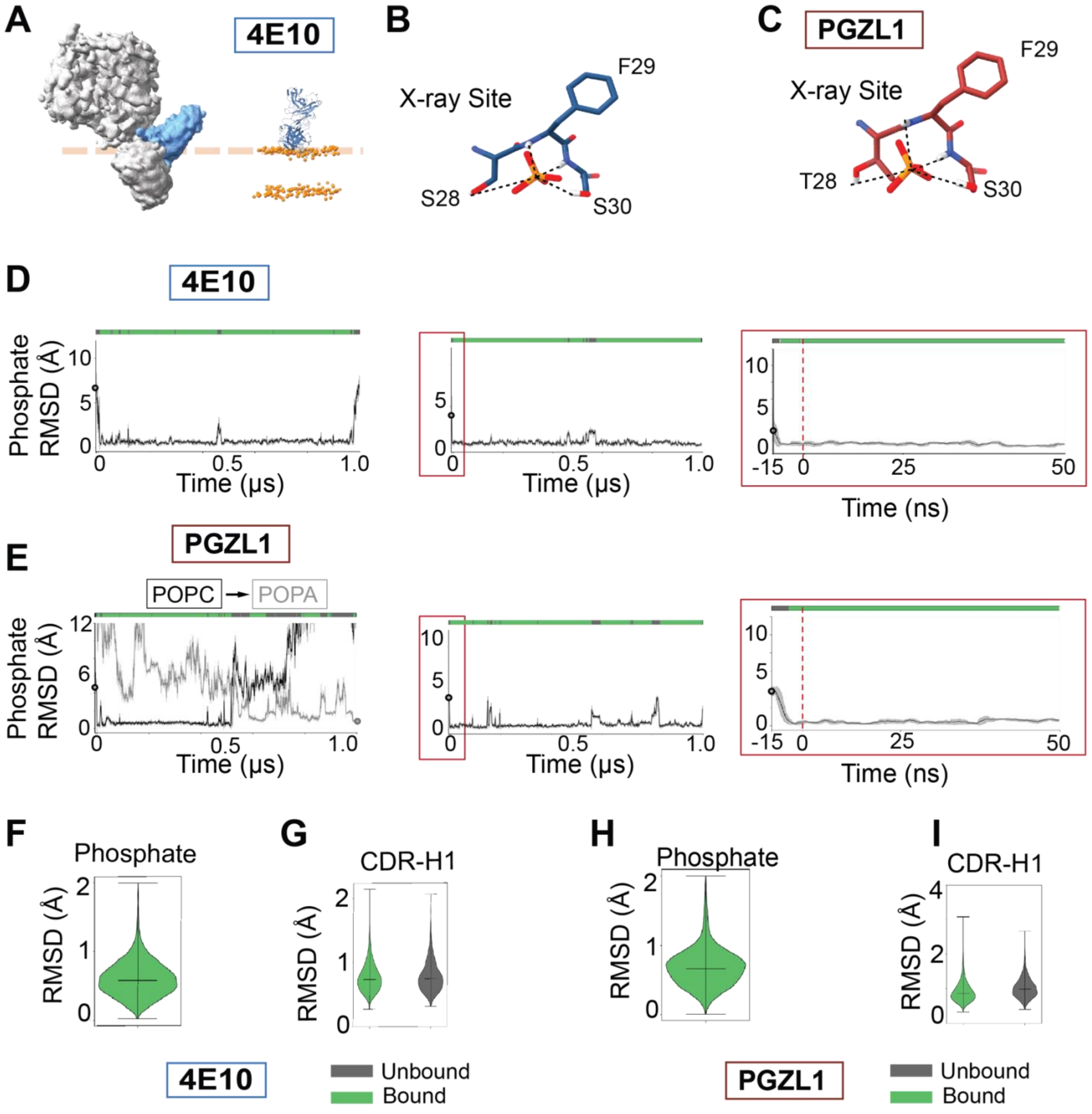

**Figure 1 Figure Supplement 1. All atom MD replicates for 4E10 and PGZL1 with phosphate group interactions**

**(A)** Theorized orientation of titled full-length HIV Env with one 4E10 Fab bound (EMD: 25024) cryoEM density map compared to a model of 4E10 Fab docked in a simplified HIV-like model membrane with PPM2.0.

**(B)** A phosphate was defined as "bound" by defining the relative position after aligning 4E10 CDR-H1 backbone conformation of each simulation frame with that of the X-ray structure: relative distance of the phosphate atom of <2.0 Å the position of the phosphate in the crystallographic loop.  As well, to control for potential backbone fluctuations, we also considered nearby positioning of at least two of the original hydrogen bond donor side chain or backbone donors within 5.25 Å to the central phosphate – accounting for the P-O bond length plus the distance of a weak hydrogen bond.

**(C)** A phosphate was defined as bound in the PGZL1 CDR-H1 loop with the same distance metrics as detailed in (**A**) for 4E10.

**(D)** Two additional replicates of 4E10 bnAb Fab all-atom MD simulations tracking reproducible lipid phosphate binding events. Right, for one replicate binding occurs as early as the preceding 15 ns NPT equilibration simulation where the protein structure is restrained but lipids can freely diffuse. The bound phospholipid-CDR complex proceeds stable through its corresponding 1 µs production simulation.

**(E)** Left, PGZL1 bnAb Fab all-atom simulation replicate showing example of a replacement event, where one lipid phosphate group binds at the known interaction site, then exchanges with the headgroup of second lipid within the course of 1 µs. Right, additional MD simulation replicate of PGZL1 where lipid phosphate is bound rapidly, during the restrained NPT stage of the simulation (middle, right).

**(F)** RMSD per frame distribution of lipid phosphate headgroup position compared to x-ray structure positions during bound time, showing sub-Å accuracy to experiment.

**(G)** RMSD per frame distribution of 4E10 CDR-H1 loop backbone coordinates versus averaged reference coordinates show the rigidity of the loop during phospholipid bound (green) and unbound (grey) time from 4 us aggregated simulation time, showing sub-Å RMSF.

**(H)** RMSD per frame distribution of PGZL1 CDR-H1 loop backbone coordinates versus averaged reference coordinates for 4 us aggregated simulation time, showing sub-Å RMSF.

**(I)** RMSD per frame distribution of lipid phosphate headgroup position compared to x-ray structure positions during bound time for PGZL1, showing sub-Å accuracy to experiment.

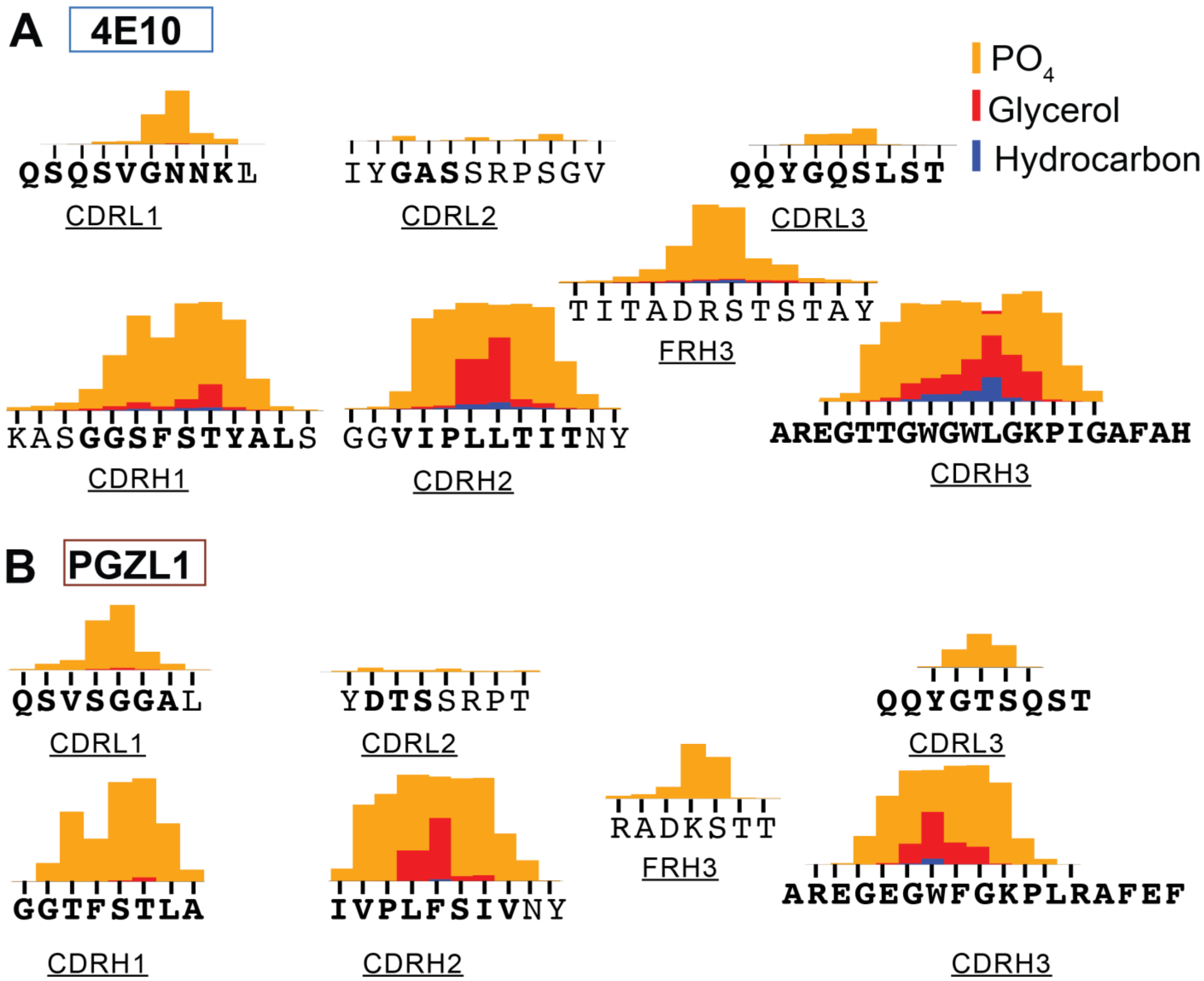

**Figure 1 Figure Supplement 2. Detailed per-residue protein-lipid interaction analysis aggregated across simulations including primary sequence to fine 4E10 and PGZL1 bnAbs membrane-interacting region**

- **(A)** Per-residue interaction profiles aligned with primary sequence for 4E10 CDR and framework regions, highlighting all residues which embed or peripherally contact lipids, defined by layers of different membrane depth and chemical environment, colored as in Figure 1E. Amino acids specifically within loops regions are bolded, with flanking membrane-contacting amino acid segments included for context.
- **(B)** Per-residue interaction profiles for lipid-interacting PGZL1 CDR loops and framework regions.

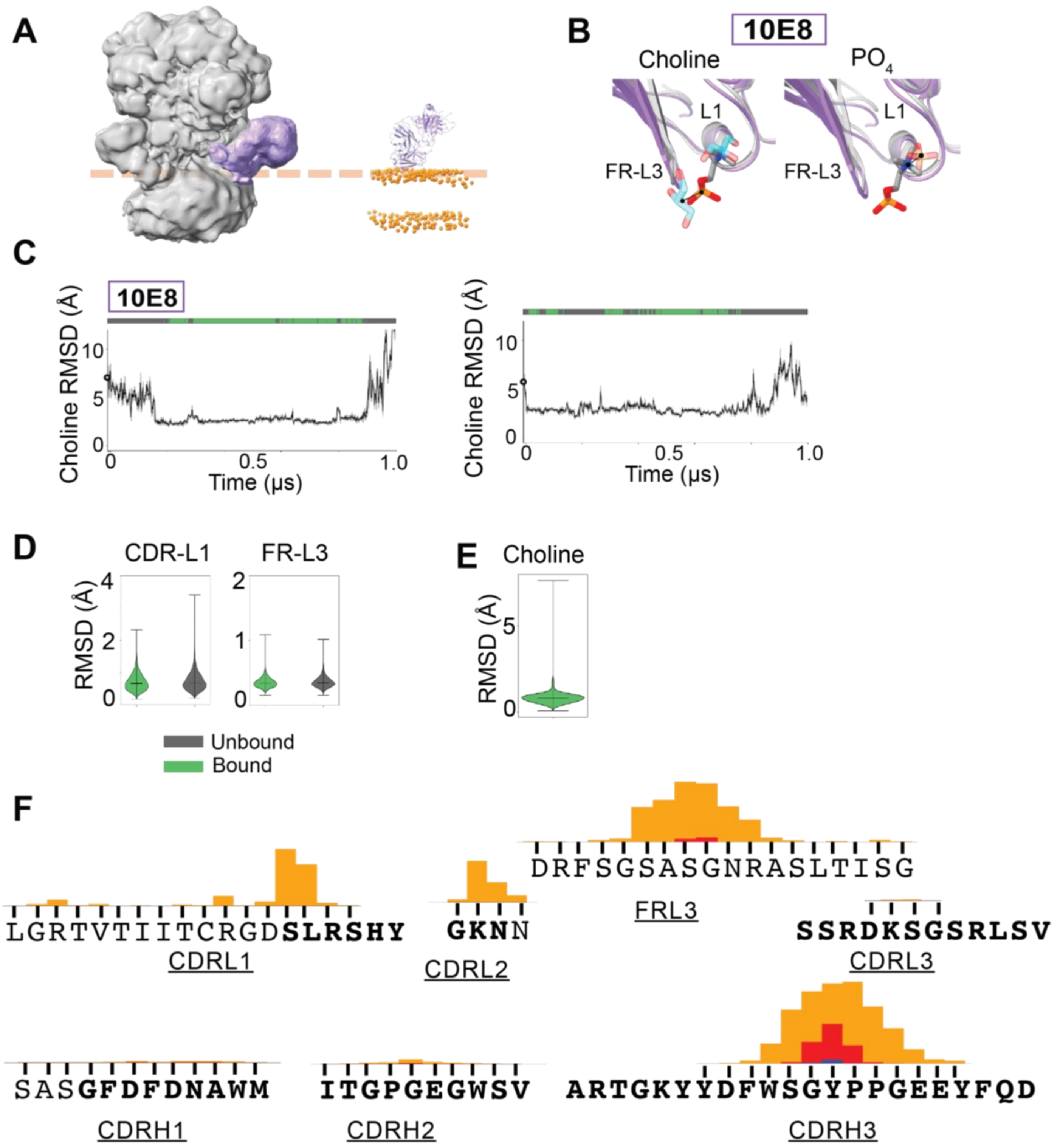

**Figure 2 Figure Supplement 1. All atom MD replicates for 10E8 detailing molecular interactions in phospholipid binding and membrane association**

**(A)** Theorized orientation of full-length HIV Env with 3 10E8 Fabs bound (EMD: 25024) cryoEM density map compared to a model of 10E8 Fab docked in a simplified HIV-like model membrane with PPM2.0.

**(B)** In the 10E8 CDR-L1 FR-L3 groove binding site, glycerols (left, partially transparent) and phospholipid headgroups (right, partially transparent) are shown overlain with the bivalent POPC headgroup bound in MD simulations (grey backbone for X-ray, purple for MD). The lipid choline cation forms complementary electrostatic interacts with the CDR-L1 unpaired backbone carbonyls in a helical loop conformation, forming a C-cap style interaction, and overlays with density modeled as free dihydrogen phosphate, lipid phosphoadic, or free glycerol in X-ray structures (Ref 24). The lipid phosphate accepts hydrogen bonds from the FR-H3. Combination of satisfied polar interaction to both lipid phosphate and choline (<3.5 Å distance) as well as choline RMSD to the relative x-ray structure positions were used to define a phospholipid as "bound" in 10E8 simulations.

**(C)** Two additional replicate all-atom MD simulations of 10E8 bnAb Fab started from varied orientations relative to the membrane, tracking RMSD of MD choline coordinates during the trajectory versus the reference phosphate coordinate from a 10E8 X-ray structure.

**(D)** RMSD per frame distribution of 10E8 CDR-L1 and FR-L3 backbone coordinates versus averaged reference coordinates for 4 us aggregated simulation time, showing sub-Å RMSF

**(E)** RMSD of lipid choline headgroup position compared to the heavy atom CDR-L1 binding site position in X-ray structures during the "bound" state for 10E8, showing sub-Å accuracy to experiment.

**(F)** Per-residue interaction profiles for lipid-interacting 10E8 CDR loops and interacting framework regions.

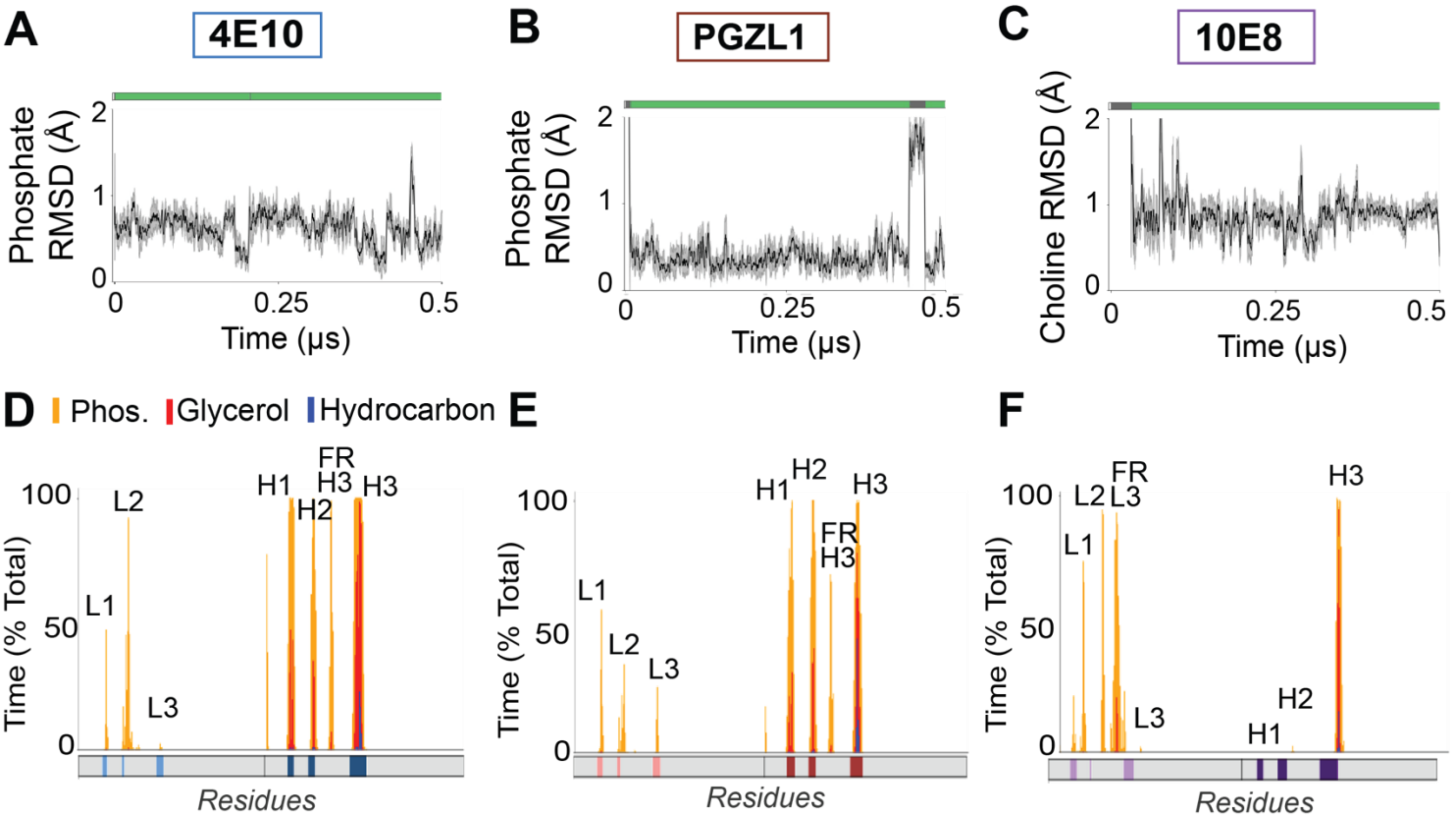

**Figure 2-Figure Supplement 2. Atomic simulations robustly capture phospholipid binding within predicted sites in realistic HIV-like membranes**

- **(A)** 500 ns simulation of 4E10 bnAb Fab all-atom MD simulations tracking a POPC lipid phosphate binding in a more realistic HIV-like membrane composition.
- **(B)** PGZL1 bnAb Fab all-atom MD simulation tracking a PSM lipid phosphate binding events in a more realistic HIV-like membrane composition.
- **(C)** 10E8 bnAb Fab all-atom MD simulations tracking choline binding in a more realistic HIV-like membrane composition.
- **(D)** Per-residue interaction profiles for lipid-interacting 4E10 CDR loops and framework regions in a realistic HIV-like lipid membrane. Interactions are colored the same as Figure 1E and reported as percent of total time from 500 ns simulations.
- **(E)** Per-residue interaction profiles for lipid-interacting PGZL1 CDR loops and framework regions in a realistic HIV-like lipid membrane.
- **(F)** Per-residue interaction profiles for lipid-interacting 10E8 CDR loops and framework regions in a realistic HIV-like lipid membrane.

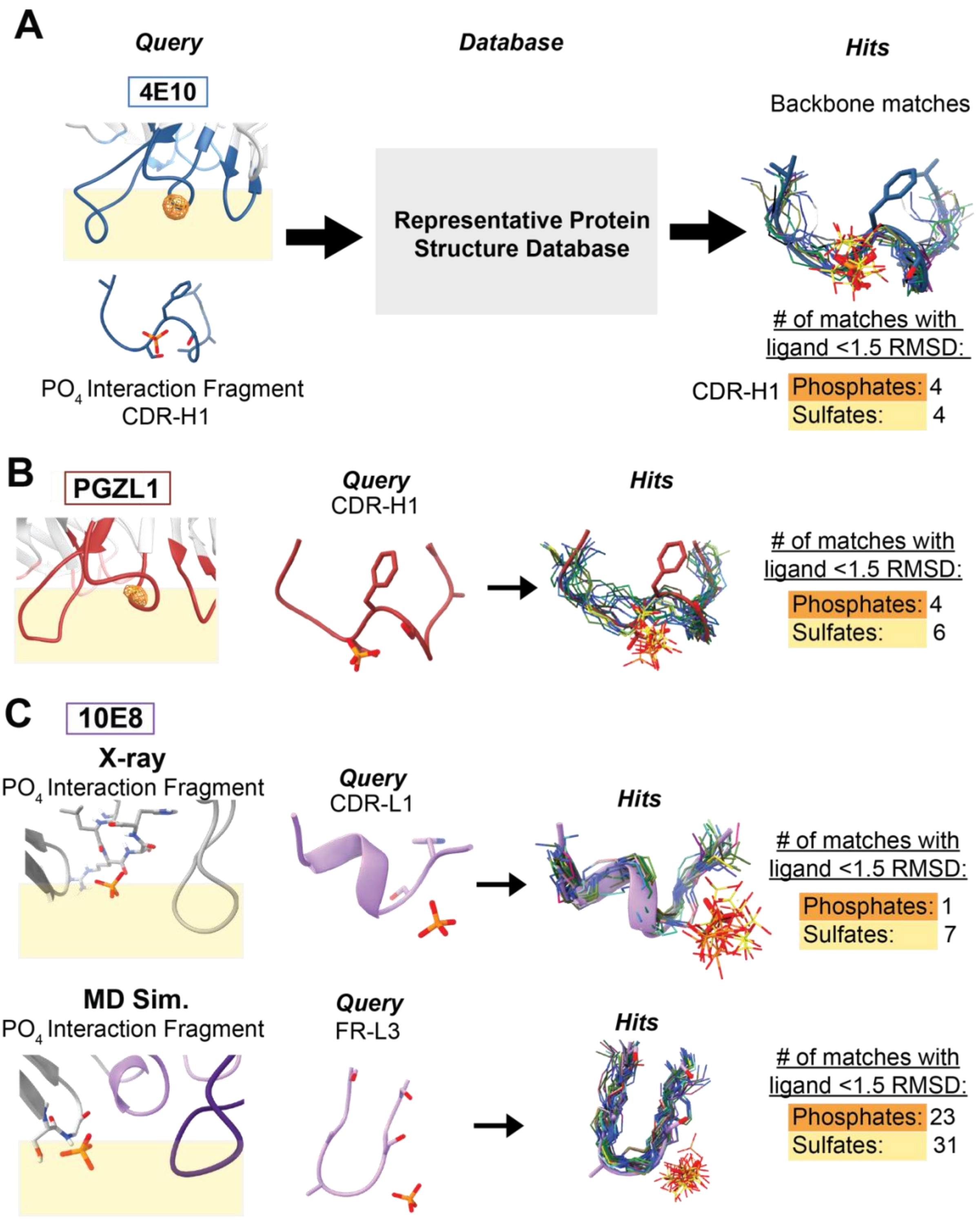

**Figure 2 Figure Supplement 3. Structural database mining shows bnAb CDR phosphate sites from MD simulations are native-like and present in the proteome.**

**(A)** Local loop geometry from each bnAb Fab simulation was extracted and queried against a 33,372 member database of experimental protein X-ray structures at < 2.5 Å resolution and <90% sequence identity previously described in Ref 37. Those protein structures from the database with loops having high structural similarity to the bnAb Fab loops (< 2 backbone atom RMSD) were identified, then further mined for the presence of crystallographic phosphate or phosphoryl groups bound in the loop within a 1.5 Å radius the reference phosphate coordinate in the experimental 4E10 bnAb structure at the CDR H1 loop. The example of the 4E10 CDR-H1 phosphate site is shown, with 4 matching examples of similar loops of distinct primary sequence binding phosphates analogously to 4E10: 3 matches from a family of bacterial bis-phosphate aldolase enzymes with organophosphate ligands, and 1 from human chymotrypsin C ($PO_4$ ligand). Many sulfate and sulfo-group

containing ligands were found as well, including the ‘positive control’ of similar PGZL1 with a $SO_4$ bound at CDRH1 (PDB:6O3K).
**(B)** Examples of loops matching PGZL1 CDR-H1 with bound phosphate/phosphoryl groups and sulfate/sulfo groups were similar to those for 4E10, in line with these two antibodies’ loops’ structural similarity.
**(C)** Given the shift of lipid phosphate site in the previous X-ray structures versus our MD simulations, we queried both the CDR-L1 loop in the X-ray structure and the FR-L2 separately to interrogate the biological relevance of each putative phosphate binding site through its prevalence in natural proteins of distinct structure. The CDR-L1 phosphate binding site (top) has a geometry which proved to be infrequent in the PDB, matching only 1 other instance. In contrast, the FR-L2 phosphate site (bottom) discovered using MD was far more common in the proteome, found in different protein families including ligase, lyase, and oxidoreductase enzymes with organic phosphoester-containing ligands.

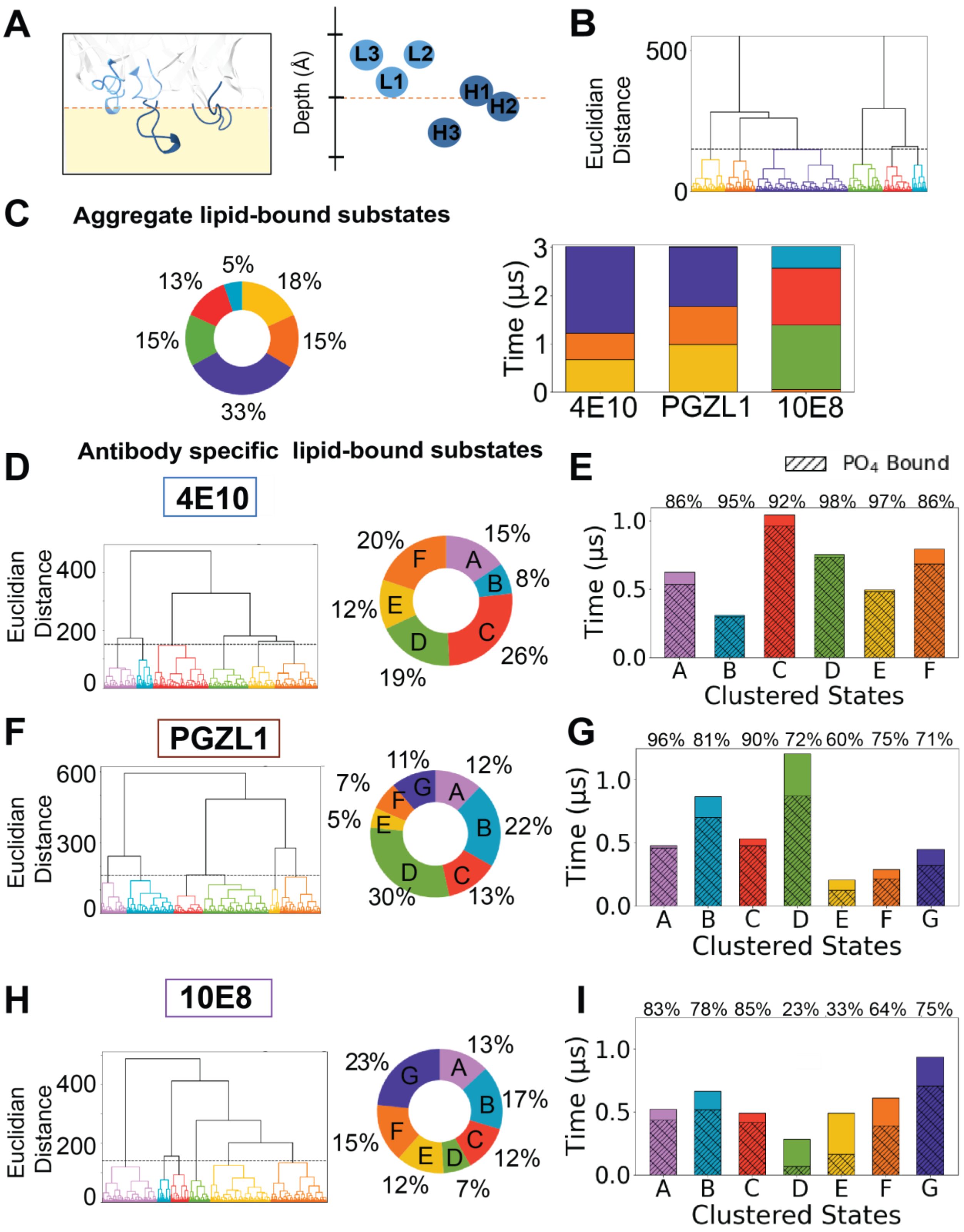

**Figure 2-Figure supplement 4. Aggregated and independent geometric clustering of bnAb surface-bound conformations**

**A)** Geometric representation of CDR loop depths in membrane where each loop is represented by the center of mass of loop residues and the top of the membrane is represented by average Z-position of lipid phosphates in upper bilayer leaflet.

**(B)** Hierarchical clustering tree of 9 µs simulation time aggregated from trajectories initiated from unique starting membrane-bound conformations (0 degrees, -15 degrees, 15 degrees initialization relative to the docked pose) for the three antibodies 4E10, PGZL1, and 10E8 structurally clustered together aggregated for their

protein-membrane interaction geometries. The features defining the substates, i.e. the feature vector used for structural clustering, include CDR loop depths as well as the Fab global approach and rotational angles (see methods).
**(C)** Classification of sub-states after aggregate clustering of each bnAb Fab's 3 μs simulation time together, clusters and percent of frames each represented by distinct colors (left). Frames from 4E10 and PGZL1 simulations cluster together, while 10E8 frames are mostly structurally divergent (right).
**(D,E,F)** Left, geometric clustering and distribution of time each Fab spent in each lipid-bound micro-state across aggregate 4 μs MD simulation time defined from independent clustering. Right, within each lipid-bound substate, the percentage of time with a phosphate group or headgroup defined as bound to the observed interaction site is listed and denoted as hatched shading on each bar graph.

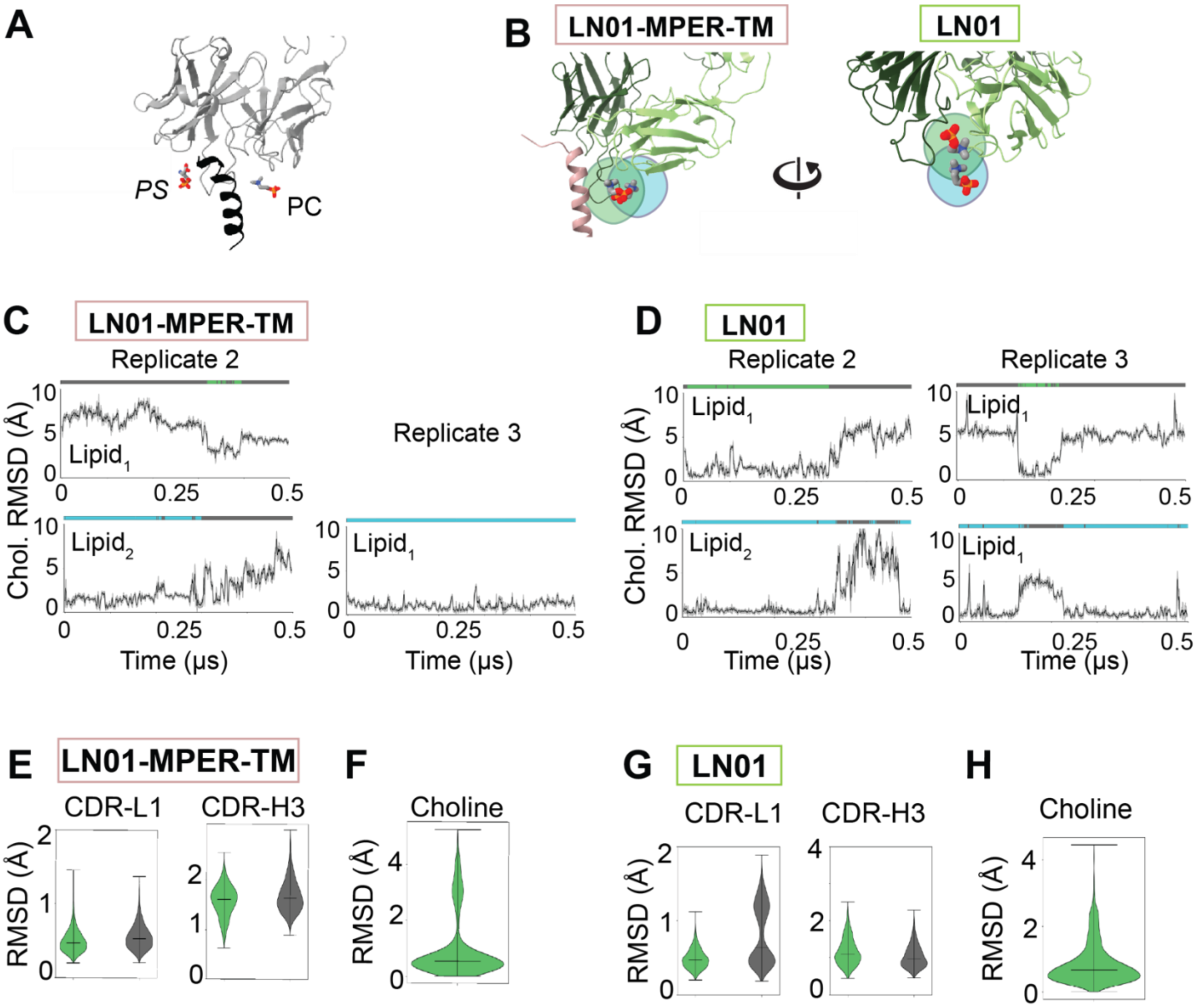

**Figure 3-Figure supplement 1. All atom MD replicates LN01-MPER-TM and LN01 with phosphate group interactions in x-ray and loading sites**

- **(A)** Composite of LN01-TM X-ray structures with phosphocholine (PC) bound within the groove of TM and CDR-H3-L1-L2 site (PDB: 6SND, 6SNE, 6SNC) and phosphatidyl-l-serine (PS) bound in a solvent exposed CDR-H1-H2-H3 site (PDB:6SND).
- **(B)** Fabs (green cartoon) depict the different lipid binding site locations predicted within MD simulations, including a site analogous to the X-ray PC site (green outline) and the new Loading site (blue outline) positions relative to each other in TM bound and apo systems.
- **(C)** Additional replicates of all-atom MD simulations for LN01+TM showing reproducible lipid phosphate binding event in both the x-ray PC site (top, green) and the loading site (bottom, cyan). No lipid phosphate interaction in x-ray PC site was captured in Replicate 3.
- **(D)** Additional replicates of all-atom MD simulations for LN0 showing reproducible lipid phosphate binding event in both the x-ray PC site (top, green) and the loading site (bottom, cyan). A lipid exchange event between x-ray and loading sites is captured in Replicate 3.
- **(E)** RMSD per frame distribution of LN01 CDR-L1 and CDR-H3 backbone coordinates versus averaged reference coordinates for LN01+MPER-TM aggregated simulation time.
- **(F)** RMSD of lipid choline headgroup position compared to x-ray structure positions during bound time for LN01+MPER-TM systems.

**(G)** RMSD per frame distribution of LN01 CDR-L1 and CDR-H3 backbone coordinates versus averaged reference coordinates for LN01 aggregated simulation time.

**(H)** RMSD of lipid choline headgroup position compared to x-ray structure positions during bound time for LN01 systems.

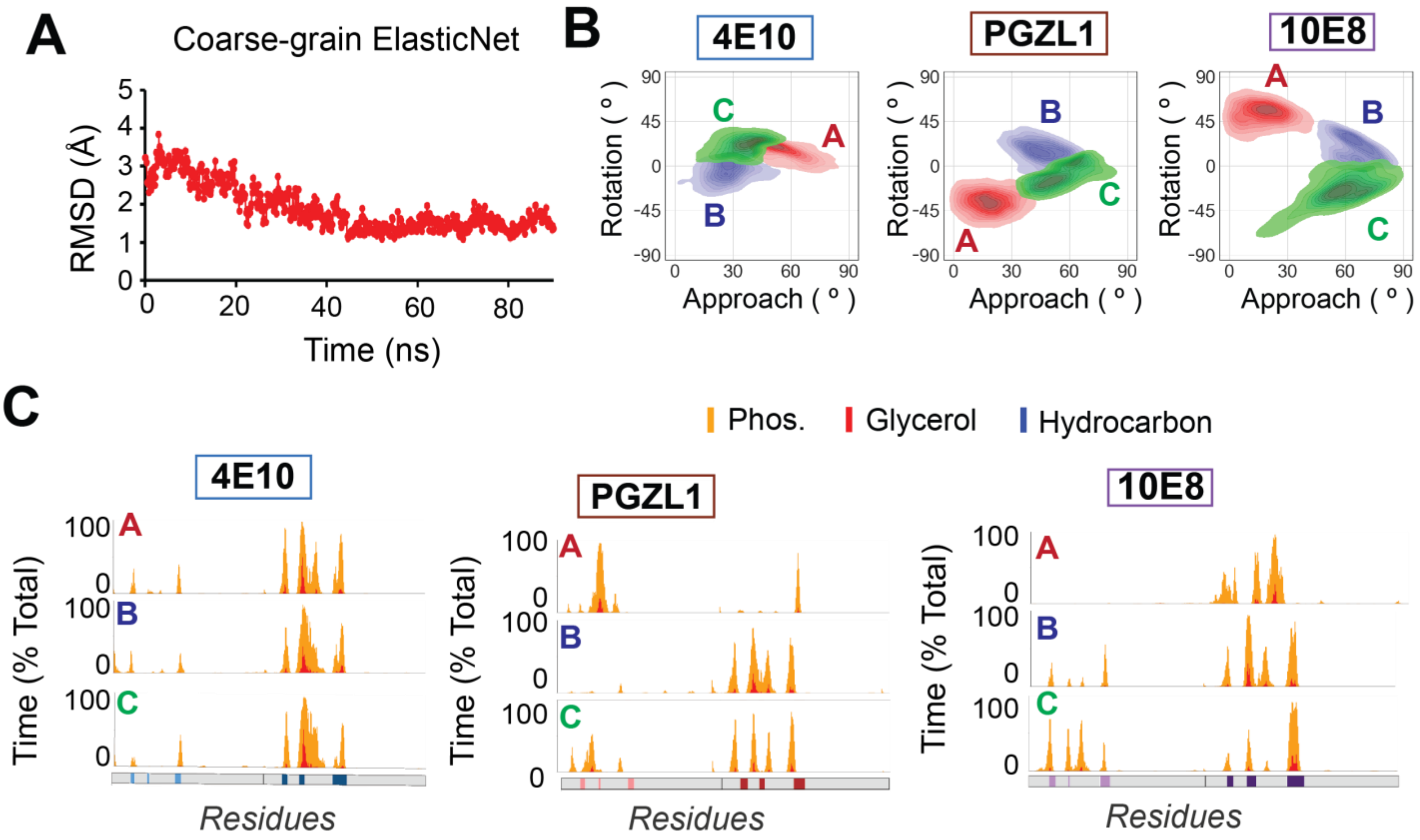

**Figure 5-Figure Supplement 1. Stability and membrane interactions from coarse-grain expanded sampling**

**(A)** The backbone RMSD of a 4E10 Fab coarse grained with the Martini force field and restrained with the ElasticNet model maintains.

**(B)** Frequency of membrane interaction angles from coarse grain spontaneous insertion as clustered by geometric substates for 4E10, PGZL1 and 10E8, colored and contoured as in Fig 3B.

**(C)** Interaction profiles for the coarse-grained spontaneously inserted frames that were clustered into substates A-C based on geometric angles. The percent of total time of each coarse grain substate for 4E10 (left), PGZL1 (middle), and 10E8 (right).

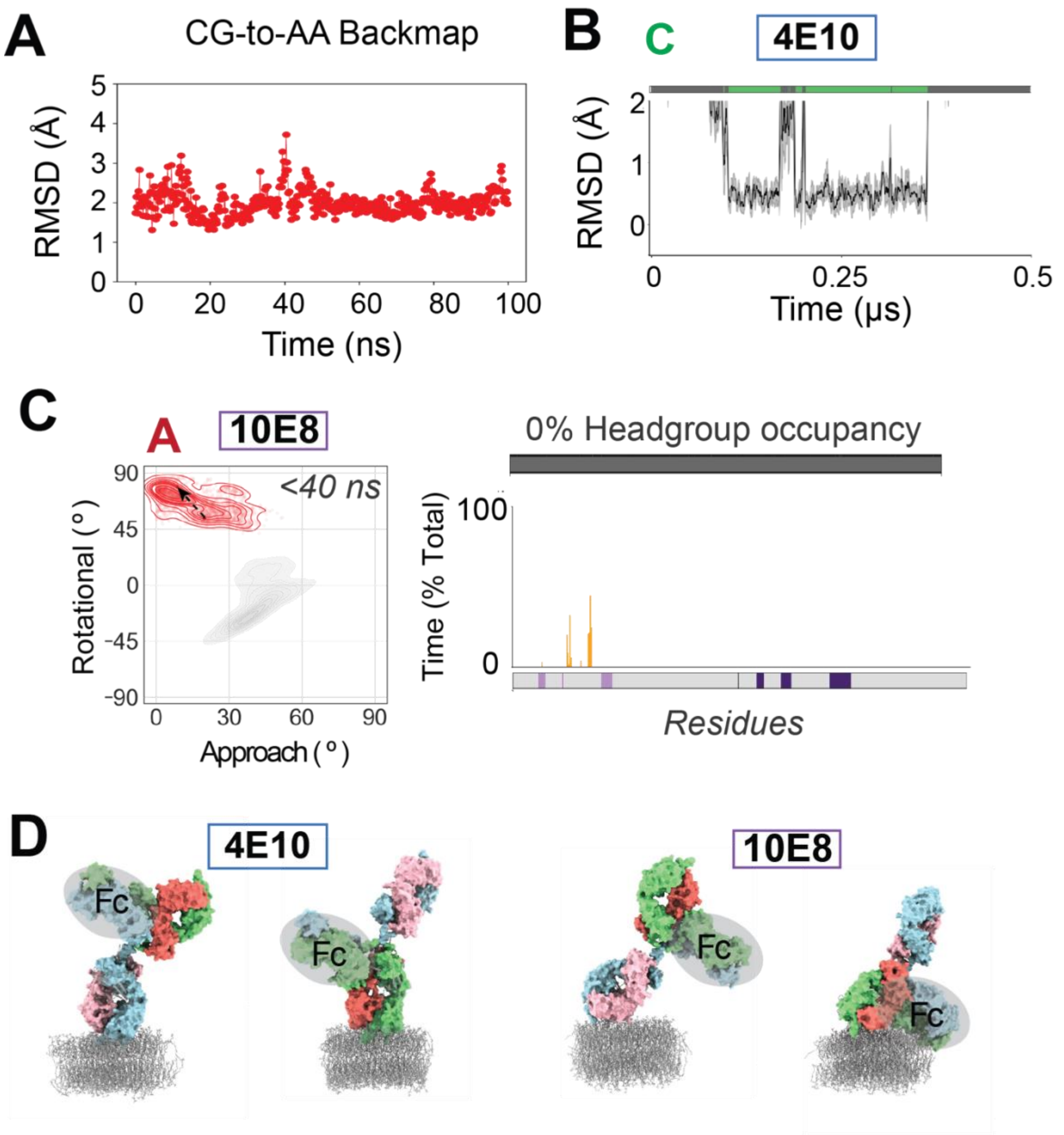

**Figure 6 Figure Supplement 1 Coarse grain to all atom backmapping validation**

- **A)** The backbone RMSD of a 4E10 Fab backmapped from coarse grain to atomistic representation remains stable in an all-atom equilibration simulation.
- **B)** The backmapped atomic State C for 4E10 demonstrated lipid headgroup binding in the expected CDR-H1 site.
- **C)** Backmapped state A for 10E8 proved an artifact after dissociating within 40 ns of atomic simulation. We observed alternative geometry of Fab domain by global orientation angles sampled (red contour) compared to our previous atomic simulations (grey contour, from Figure 2), no lipid headgroup binding (right, top), and a distinct protein-lipid interaction profile during the contact time with sparse surface interactions with lipids (right, bottom).
- **D)** Potential for bivalent Fab domain insertion for the full length IgG of 4E10 (left) based on the membrane-bound conformation, built by structural alignment to a full-length IgG model (PDB: 1HZH). Simultaneous Fab contact from both IgG arms appears improbable based on strain and rotational rearrangements that would be imposed to the IgG hinge region to accommodate such a pose. Frames extracted from 10E8 simulations (right) where a full-length IgG model (PDB: 1HZH) is superposed to membrane-bound Fab imply unlikely bivalent interactions. Representative frames are from 2 distinct time points from 0-degree tilt replicates where phospholipids were bound for both 4E10 and 10E8,

representing at least 2 distinct macroscopic substates differing in global light chain and heavy chain orientation towards the membrane between the two antibodies.

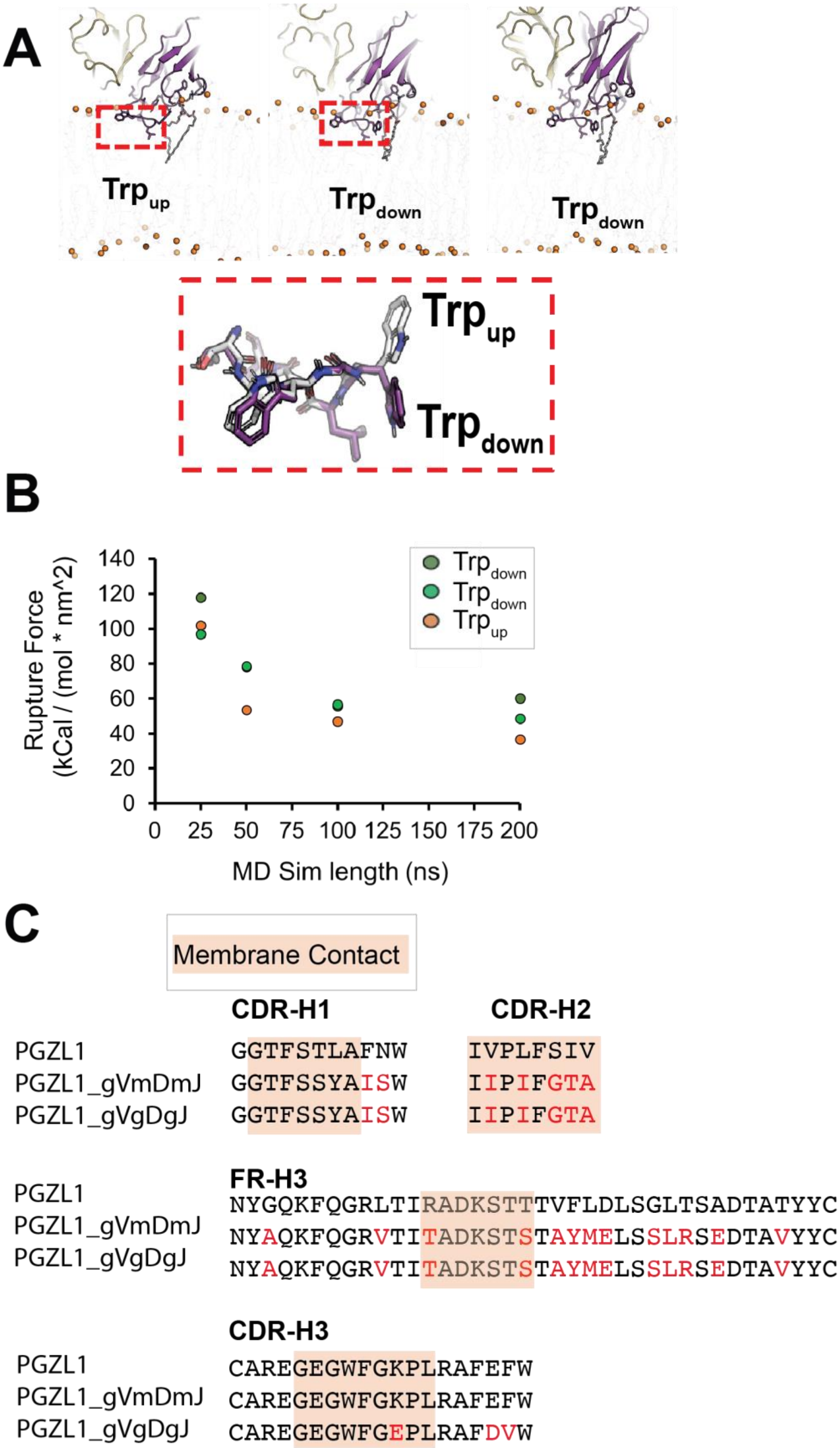

**Figure 7 Figure Supplement 1. Constant velocity pulling simulations are sensitive to dynamics and sequence variation**

**(A)** Rotameric states of Trp100a and Trp100b residues in the CDR-H3 loop of 4E10. Distinct starting conformations of TrpUp and TrpDown models used in pulling simulations are shown.

**(B)** Max force for each wildtype 4E10 biased membrane-dissociation replicate simulation is plotted against the time of pulling simulation across fixed distance from the membrane, thus testing different constant pulling velocities. A clear difference for rupture force is detectable for 4E10 replicates initiated from the TrpUp versus TrpDown conformation, most noticeable at 50, 100, and 200 ns. Pulling trajectories are nearly identical for replicates initiated from poses adopting two TrpDown state (green) at 50 and 100 ns. Max force values roughly plateau between 100 and 200 ns.

**(C)** A sequence alignment for germline, intermediate, and mature PGZL1 variants of regions that contact the membrane during primary atomistic simulations with mature PGZL1 (CDR-H1,H2,H3 and FR-H3).

**Supplementary Table 1. Bioinformatics descriptions for ligand bound structural matches**
Lipid interacting loop fragments from MD simulations or crystal structures were queried against structural database and hits were filtered to include ligands within 1.5 angstroms from experimental or simulation predicted phosphate or choline interactions. Relevant information for hits from each queried loop fragment includes PDB code, structural entry name in The Protein Data Bank, bound ligand ID, organism from which the protein structure is isolated, and functional classification.

| PDB ID | bnAb | Ligand | Organism | Classification |
|---|---|---|---|---|
| 3LUL | 10e8 | LLP | legionella pneumophila | lyase |
| 6RSW | 10e8 | EPE | oryctolagus cuniculus | contractile protein |
| 2F99 | 10e8 | SO4 | streptomyces galilaeus | biosynthetic protein |
| 3JZ6 | 10e8 | PLP | mycobacterium smegmatis | transferase |
| 7LV7 | 10e8 | LLP | giardia intestinalis | transferase |
| 3I3L | 10e8 | FAD | streptomyces venezuelae | hydrolase |
| 5BVA | 10e8 | FAD | streptomyces | hydrolase |
| 4UU9 | 10e8 | SO4 | homo sapiens | immune system |
| 3A0E | 10e8 | SO4 | polygonatum cyrtonema | sugar binding protein |
| 6R5M | 10e8 | SO4 | dendroaspis polylepis | toxin |
| 5BUK | 10e8 | FAD | streptomyces sp. cnq-418 | oxidoreductase |
| 5UAO | 10e8 | FAD | microbispora sp. atcc pta-5024 | oxidoreductase |
| 1OCY | 10e8 | SO4 | bacteriophage t4 | structural protein |
| 6T21 | 10e8 | 5CM | escherichia coli (strain k12) | hydrolase |
| 5TUI | 10e8 | SO4 | uncultured bacterium | oxidoreductase |
| 6K8V | 10e8 | SO4 | porphyrobacter dokdonensis dsw-74 | oxidoreductase |
| 6JIF | 10e8 | PLP | pseudomonas sp. uw4 | transferase |
| 1O57 | 10e8 | SO4 | bacillus subtilis | dna binding protein |
| 6XGS | 10e8 | PO4 | brucella suis biovar 1 | biosynthetic protein, lyase |
| 3GG8 | 10e8 | SO4 | toxoplasma gondii | transferase |
| 3WUD | 10e8 | SO4 | xenopus laevis | sugar binding protein |
| 4TVI | 10e8 | LLP | brucella abortus | transferase |
| 2Y4R | 10e8 | PLP | pseudomonas aeruginosa | lyase |
| 2YIL | 10e8 | SO4 | sarcocystis muris | sugar binding protein |
| 3CW9 | 10e8 | AMP | alcaligenes sp. | ligase |
| 2OVR | 10e8 | SO4 | homo sapiens | transcription/cell cycle |
| 3UGF | 10e8 | SO4 | pachysandra terminalis | transferase |
| 5IOB | 10e8 | MES | corynebacterium glutamicum (strain atcc 13032 /dsm 20300 / jcm 1318 / lmg 3730 / ncimb 10025) | hydrolase |
| 1HM6 | 10e8 | SO4 | sus scrofa | metal, lipid binding protein |
| 3CSW | 10e8 | PLP | thermotoga maritima msb8 | transferase |
| 6Q1S | 10e8 | PMP | mycobacterium tuberculosis (strain atcc 25618 /h37rv) | transferase |
| 1M0W | 10e8 | SO4 | saccharomyces cerevisiae | ligase |
| 6UL2 | 10e8 | SO4 | uncultured bacterium | flavoprotein |
| 6NSD | 10e8 | FAD | saccharomonospora sp. cnq490 | biosynthetic protein |

| | | | | |
|---|---|---|---|---|
| 4NZF | 10e8 | SO4 | geobacillus stearothermophilus | hydrolase |
| 4PZ7 | 10e8 | SO4 | schizosaccharomyces pombe | transferase |
| 5WGX | 10e8 | FAD | malbranchea aurantiaca | oxidoreductase |
| 1I2K | 10e8 | PLP | escherichia coli | lyase |
| 5EWO | 10e8 | SO4 | human astrovirus-1 | viral protein |
| 2YDT | 10e8 | SO4 | gibberella zeae | hydrolase |
| 6NST | 10e8 | SO4 | pseudomonas aeruginosa (strain atcc 15692 / dsm22644 / cip 104116 / jcm 14847 / lmg 12228 / 1c / prs 101 / pao1) | transferase |
| 4PQQ | 10e8 | PO4 | mus musculus | protein binding |
| 2XIW | 10e8 | SO4 | sulfolobus acidocaldarius | dna binding protein |
| 4G1Q | 10e8 | SO4 | human immunodeficiency virus type 1 | transferase, hydrolase |
| 3M4F | 10e8 | CXS | scytalidium acidophilum | hydrolase |
| 5MP6 | 10e8 | SO4 | homo sapiens | immune system |
| 6BB9 | 10e8 | MES | salmonella typhimurium | lyase |
| 2AQJ | 10e8 | FAD | pseudomonas fluorescens | biosynthetic protein |
| 4CU7 | 10e8 | SO4 | streptococcus pneumoniae tigr4 | hydrolase |
| 4K6N | 10e8 | PLP | saccharomyces cerevisiae | lyase |
| 6Q0S | 10e8 | SO4 | human respiratory syncytial virus (subgroup b /strain 18537) | immune system |
| 6UI5 | 10e8 | FAD | streptomyces sp. nrrl 11266 | oxidoreductase |
| 3NYQ | 10e8 | AMP | streptomyces coelicolor | ligase |
| 2GFQ | 10e8 | SO4 | pyrococcus horikoshii | unknown function |
| 3KDW | 10e8 | PO4 | bacteroides vulgatus atcc 8482 | sugar binding protein |
| 3FP3 | 10e8 | SO4 | saccharomyces cerevisiae | transport protein |
| 4I42 | 10e8 | SO4 | escherichia coli | lyase |
| 3EUP | 10e8 | SO4 | cytophaga hutchinsonii | transcription regulator |
| 1UQR | 10e8 | SO4 | actinobacillus pleuropneumoniae | lyase |
| 2HYJ | 10e8 | SO4 | streptomyces coelicolor | transcription |
| 6QOK | 10e8 | SO4 | mycobacterium abscessus | transferase |
| 5AVM | 10e8 | SO4 | thermus thermophilus hb8 | ligase |
| 5MQQ | 4E10 | SO4 | corynebacterium glutamicum | transcription |
| 3QM3 | 4E10 | SO4 | campylobacter jejuni | lyase |
| 3Q94 | pgzl1 | 13P | bacillus anthracis | lyase |
| 3GN3 | pgzl1 | SO4 | pseudomonas syringae pv. tomato | unknown function |
| 1GVF | pgzl1 | PGH | escherichia coli | lyase |
| 2AIB | pgzl1 | MES | phytophthora cinnamomi | toxin |
| 4H4F | pgzl1 | PO4 | homo sapiens | hydrolase |
| 7NC7 | pgzl1 | 13P | bacillus methanolicus (strain mga3 / atcc53907) | lyase |
| 1CR1 | pgzl1 | SO4 | enterobacteria phage t7 | transferase |
| 5MIY | pgzl1 | SO4 | legionella pneumophila | ligase |
| 6QE8 | pgzl1 | SO4 | aspergillus niger | hydrolase |
| 5NN4 | pgzl1 | SO4 | homo sapiens | hydrolase |

**Supplementary Table 2. Detailed definitions of substates from global clustering of atomic geometries** Statistics of each metric used to define geometric macro-substates in global clustering which included aggreagating all simulation time from 10E8, PGZL1, and 4E10 together (Figure 2-figure supplement 1A-C). Mean and standard deviation for CDRL loop depths, CDRH loops depths and angles of approach and rotation extracted from each substate are reported.

| | **CDRL1 Depth** | | | | **CDRH1 Depth** | |
|---|---|---|---|---|---|---|
| **Cluster** | **Mean** | **Standard Deviation** | | **Cluster** | **Mean** | **Standard Deviation** |
| Yellow | 10.37 | 2.72 | | Yellow | 3.28 | 2.28 |
| Orange | 8.35 | 3.12 | | Orange | 3.96 | 2.21 |
| Purple | 7.24 | 2.29 | | Purple | 3.02 | 2.04 |
| Green | 8.47 | 1.44 | | Green | 29.81 | 3.12 |
| Red | 7.8 | 1.67 | | Red | 21.33 | 2.72 |
| Cyan | 11.41 | 2.32 | | Cyan | 11.57 | 3.55 |
| | **CDRL2 Depth** | | | | **CDRH2 Depth** | |
| **Cluster** | **Mean** | **Standard Deviation** | | **Cluster** | **Mean** | **Standard Deviation** |
| Yellow | 12.27 | 2.55 | | Yellow | 1.1 | 1.7 |
| Orange | 7.11 | 2.5 | | Orange | 5 | 1.56 |
| Purple | 8.3 | 1.9 | | Purple | 2.38 | 1.54 |
| Green | 7.72 | 1.94 | | Green | 27.77 | 2.31 |
| Red | 5.16 | 1.56 | | Red | 20.71 | 2.14 |
| Cyan | 5.77 | 2.02 | | Cyan | 13.51 | 3.91 |
| | **CDRL3 Depth** | | | | **CDRH3 Depth** | |
| **Cluster** | **Mean** | **Standard Deviation** | | **Cluster** | **Mean** | **Standard Deviation** |
| Yellow | 11.94 | 2.31 | | Yellow | 5.96 | 2.49 |
| Orange | 12.05 | 2.31 | | Orange | 3.54 | 1.3 |
| Purple | 10.34 | 2 | | Purple | 3.95 | 1.38 |
| Green | 16.48 | 1.58 | | Green | 15.31 | 2.04 |
| Red | 15.29 | 1.67 | | Red | 10.74 | 1.55 |
| Cyan | 16.65 | 2.03 | | Cyan | 8.57 | 1.6 |
| | **Approach Angle** | | | | **Rotation Angle** | |
| **Cluster** | **Mean** | **Standard Deviation** | | **Cluster** | **Mean** | **Standard Deviation** |
| Yellow | 71.24 | 5.26 | | Yellow | 9.42 | 5.64 |
| Orange | 67.25 | 5.22 | | Orange | -13.31 | 5.79 |
| Purple | 80.2 | 4.16 | | Purple | -2.86 | 5.06 |
| Green | 34.96 | 5.23 | | Green | -29.46 | 5.66 |
| Red | 46.9 | 5.41 | | Red | -9.83 | 5.55 |
| Cyan | 43.41 | 6.17 | | Cyan | 11.27 | 10.22 |

**Supplementary Table 3. Spontaneously inserted coarse grain membrane contact events for bnAb Fabs**
Description of 14 μs coarse grain simulations for spontaneously inserting 4E10, PGZl1, and 10E8 antibody Fabs. Description of 10 μs coarse grain simulations for negative control systems using BSA and 13h11 Fab are included. Membrane association time is reported for each replicate that was initiated from various orientations relative to the membrane plane, along with total membrane contact time for each Fab system. Average contact time and standard deviation from total membrane contact time is reported. Average contact time and standard deviation per each association event in each antibody systems is also reported.

| **Antibody/ *Replicate*** | **Membrane Contact Time (μs)** | | **Membrane Contact Time (μs)** | | **Membrane Contact Time (μs)** | | **Membrane Contact Time (μs)** | | **Membrane Contact Time (μs)** |
|---|---|---|---|---|---|---|---|---|---|
| **4E10** | | **PGZL1** | | **10E8** | | **BSA** | | **13h11** | |
| *01* | 7.38 | *01* | 10.65 | *01* | 9.75 | *01* | 0.06 | *01* | 0.33 |
| *02* | 1.62 | *02* | 12.27 | *02* | 3.66 | *02* | 1.71 | *02* | 0.12 |
| *03* | 0.87 | *03* | 10.53 | *03* | 0.03 | *03* | 0 | *03* | 0 |
| *04* | 9.54 | *04* | 11.88 | *04* | 11.82 | *04* | 0.09 | *04* | 0.03 |
| *05* | 0.87 | *05* | 7.47 | *05* | 0.03 | *05* | 0 | *05* | 0.03 |
| *06* | 11.82 | *06* | 9 | *06* | 11.07 | *06* | 0.06 | *06* | 0.24 |
| *07* | 0.09 | *07* | 9.66 | *07* | 11.13 | *07* | 0 | *07* | 0.06 |
| *08* | 2.94 | *08* | 0.06 | *08* | 12.42 | *08* | 0.03 | *08* | 0.27 |
| *09* | 0.12 | *09* | 5.55 | *09* | 5.16 | *09* | 0 | *09* | 0.33 |
| *10* | 0.06 | *10* | 0 | *10* | 5.58 | *10* | 0.03 | *10* | 0 |
| *11* | 9.54 | *11* | 5.04 | *11* | 11.88 | *11* | 0 | *11* | 0.54 |
| *12* | 12 | *12* | 0 | *12* | 0.09 | *12* | 0.06 | *12* | 0.15 |
| *13* | 6.24 | *13* | 2.76 | *13* | 0.06 | *13* | 0.06 | *13* | 0.03 |
| *14* | 0 | *14* | 10.62 | *14* | 0 | *14* | 0.06 | *14* | 0.27 |
| *15* | 12.93 | *15* | 8.4 | *15* | 1.14 | *15* | 3.78 | *15* | 0.03 |
| *16* | 0.21 | *16* | 12.48 | *16* | 2.28 | *16* | 0.06 | *16* | 0 |
| *17* | 11.31 | *17* | 11.4 | *17* | 0.48 | *17* | 0.12 | *17* | 0.03 |
| *18* | 0.12 | *18* | 0.09 | *18* | 3.48 | *18* | 0.09 | *18* | 0.12 |
| **Total (μs)** | 87.66 | | 127.86 | | 90.06 | | 6.18 | | 2.58 |
| **Average Aggregated Time ± S.D.** | 4.87±5.08 | | 7.10±4.67 | | 5.0±4.94 | | 0.35±0.94 | | 0.14±0.15 |
| **Average Association Event Time ± S.D** | 2.87±4.18 | | 1.36±2.0 | | 1.3±1.41 | | 0.57±0.91 | | 0.21±07 |
| **Total Time Sampled (μs)** | 252 | | 252 | | 252 | | 180 | | 180 |

**Supplementary Table 4. Benchmarking Loop Conformations**
Number of fragments found based on a range of RMSD cutoffs for structural searches with phosphate binding fragments of MPER bnAbs as input query structures. Intermediate stringency is found at a 2Å RMSD cutoff. Identified hits with a 2Å RMSD cutoff using MPER bnAb phosphate binding fragments as queries compared to other structural queries with similar length but distinct topologies.

| **RSMD Cutoff (Å)** | **Hits** | **RSMD Cutoff (Å)** | **Hits** |
|---|---|---|---|
| **4e10_CDRH1** | | **10e8_FRL3** | |
| 3 | 3103691 | 3 | 3675122 |
| 2.5 | 665906 | 2.5 | 1595534 |
| 2 | 53489 | 2 | 504500 |
| 1.5 | 3906 | 1.5 | 129694 |
| 1 | 864 | 1 | 39569 |
| 0.5 | 2 | 0.5 | 2 |
| **Query Structure** | **RSMD Cutoff (Å)** | **Hits** | **Topology** |
| 10e8_FRL3 | 2 | 504500 | 7 aa loop (beta-loop-beta) |
| 3U1S_FRL3 | 2 | 530769 | PGT145 Ab FRL3 loop (7 aa between 2 beta sheets) |
| 8ABP_A_10-16 | 2 | 1122431 | 7 AA loop beta-loop-helix |
| 5MEB_B_543-549 | 2 | 504898 | 7 AA loop helix- loop -helix |
| 4e10_CDRH1 | 2 | 53489 | 10 aa loop (beta-loop-beta) |
| 3U1S_CDRH1 | 2 | 154014 | PGT145 Ab cdrh1 loop (10 aa between 2 beta sheets) |
| 2QTF_A_332-341 | 2 | 11359 | 10 aa loop (beta-loop-helix) |
| 4LLS_B_159_168 | 2 | 187764 | 10 aa loop (helix-loop-helix) |

**Video 1 : *De novo* predicted phosphate binding in atomistic 4E10 Fab MD simulation**
Atomistic simulation of 4E10 Fab (heavy chain: dark blue, light chain: light blue) initially docked to the membrane using OPM PPM server prediction. Lipids and cholesterol are shown as grey sticks with phosphates from top and bottom leaflets shown as orange spheres. The binding phosphate (large orange sphere) from a POPC lipid is initially more than 10 Å away from CDR-H1 loop and finds interaction with CDR-H1 residues in first 200 ns of simulation and maintains for remainder of 1 µs.

**Video 2 : Phosphate binding and replacement event in PGZL1 bnAb Fab all-atom simulation**
Atomistic simulation of PGZL1 Fab (heavy chain: salmon, light chain: light pink) initially docked to the membrane using OPM PPM server prediction. Lipids and cholesterol are shown as grey sticks with phosphates from top and bottom leaflets shown as orange spheres. The first interacting phosphate from a POPC lipid (large orange sphere) is initially more than 6 Å away from CDR-H1 loop and binds in CDR-H1 loop for first 500 ns of simulation. A second phosphate from a POPA lipid (pink sphere) replaces initial interaction around 550 ns and maintains the CDR-H1 interactions for the rest of the microsecond simulation.

**Video 3 : Unbiased spontaneous membrane insertion event in coarse grain 4E10 Fab MD simulation**
Coarse grain 4E10 Fab (blue) is initialized from 2 nm above an assembled membrane (grey lipid tails and cholesterol and orange phosphates). Random diffusion and tumbling in explicit water is observed before initial membrane contact is made. Membrane association is followed by reorganization of the Fab-membrane conformation for the remainder of the 14 µs simulation.

**Data availability**

Due to file size limitations, a representative trajectory and corresponding coordinate file are publicly available for each all-atom simulation at https://zenodo.org/records/13830877. All other simulations are available by request to authors. Custom python scripts written to analyze molecular dynamic simulations are available on GitHub: https://github.com/cmaillie98/mper_bnAbs.git.

### Methods

#### Atomistic Simulations

The coordinates for Fab models were obtained from x-ray crystallography structures for each antibody (PDB id: 2FX9, 6O3D, 5T85 for 4E10, PGZL1, and 10E8 respectively). PDB 6SNE was used to generate LN01+MPER-TM systems. LN01 systems used PDB 6SND as a starting model. Any missing residues in atomic models were built with ModLoop. Original membrane docked orientations for each Fab in the bilayer were generated with Orientations of Proteins in Membranes (OPM) Positioning of Proteins in Membrane 2.0 (PPM) using per residue predicted transfer free energies from water to membrane environments. For LN01+MPER-TM systems specifically, the PPM2.0 prediction used the entire protein complex for global docking optimization. Replicates with varied membrane docked starting orientations were generated by rotating the Fab along the first principal axis of the Fab +/- 15 degrees relative to the X axis.

Using the CHARMM-GUI webserver, the pre-positioned Fab was modified to include acetylated N-terminus (ACE) and methylamidated C-terminus (CT3) capping on each chain with appropriate disulfide bonds for a typical Fab structure. The Fab was then docked in an HIV-like lipid bilayer composed of 70% palmitoyl-oleoyl phosphatidylcholine (POPC), 25% cholesterol (CHOL), and 5% palmitoyl-oleoyl phosphatidic acid (POPA) bilayer and hydrated (minimum water height of 25Å). Ions were added to neutralize and bring the system to a final concentrations of 0.15 mM KCl. Atomistic simulations were performed with Gromacs 2021 and CHARMM36 forcefield. Water molecules were described with the TIP3 model. All systems were minimized, with 5000 steps with steepest descent algorithm. A 2 fs time step was used along with the LINCS constraint algorithm during the equilibration stage. Electrostatics were treated with Particle Mesh Ewald, and the cutoff for both Coulomb and van der Waals interactions (Lennard Jones potential) was 1.2 nm. An 100 ps NVT equilibration phase applied a 4000 kJ/mol/nm$^3$ restraint on heavy atoms in protein and 1000 kJ/mol/nm$^3$ restraint on lipids with the velocity rescaling temperature coupling for protein, lipid, and solvent groups independently coupled to a 310 K bath with a temperature time constant of 0.1 ps$^{-1}$. A subsequence 15 ns NPT equilibration phase was performed with lipid restraints removed and 1000 kJ/mol/nm$^3$ restraints on the alpha carbons of protein. The Berendsen thermostat was used, with protein, lipid, and solvent independently coupled to a 310 K bath with temperature time constant 1.0 ps$^{-1}$. The Berendsen barostat was used with semi-isotropic coupling. Restraints were then removed for 0.5-1 us production run with Noose-Hoover thermostat applied to protein, lipid, and solvent groups independently coupled to a 310 K bath with temperature time constant 1.0 ps$^{-1}$. The Parrrinello-Rahman barostat was used with semi-isotropic coupling and a 5.0 ps$^{-1}$ pressure time constant .

#### Generating coarse grain models

X-ray structures were used as starting modes for each antibody (PDB id: 2FX9, 6O3D, 5T85 for 4E10, PGZL1, and 10E8 respectively). Coarse grain representations of Fab models were generated based on the Martini version 2.2 four to one mapping of heavy atoms to bead representation using the martinize script with an elastic network that defined harmonic bonds at default parameters. All coarse grain simulations were performed with Gromacs 2021 and CHARMM36 forcefield.

#### Pre-embedded coarse grain simulations

Coarse grain Fab models were varied in their pre-embedded orientation by a rotation matrix of every 90 degrees in X axis, every 90 degrees in the Y axis, and every 1 nm along Z axis for 3 nm. A total of 40 Fab orientations that were physiologically plausible and did not include the constant region deeply embedded in the membrane were placed in a 15x15x20 nm box. After adding a Fab model to a box, Martini coarse grain lipids were added to the bottom half of the box at final percentage of 70% POPC, 25% cholesterol, and 5% POPA. Ions were added to neutralize the system to a final concentration of 0.15 mM NaCl and coarse grain water molecules were added to solvate the system with an adjusted van der Waals radius of 0.21 nm. A 30 fs time step was used along with the LINCS constraint algorithm. The system was minimized for 5000 with steepest descent algorithm. Minimization was followed by a short 30 ns membrane assembly step where the cutoff for both Coulomb and van der Waals interactions (Lennard Jones potential) was 1.1 nm. The velocity rescaling thermostat was used, with protein, lipid, and solvent independently coupled to a 310 K bath with 1.0 $ps^{-1}$ temperature time constant. The Berendsen barostat was used with isotropic coupling and a 6.0 $ps^{-1}$ pressure time constant . A 5 us production simulation was then performed with velocity rescaling thermostat applied to protein, lipid, and solvent groups independently coupled to a 310 K bath with temperature time constant 1.0 $ps^{-1}$. The Parrrinello-Rahman barostat was used with semi-isotropic coupling and 12.0 $ps^{-1}$ pressure time constant.

**Spontaneously associating coarse grain simulations**

We used the same starting models and rotation matrix as in pre-embedded systems, but adjusted Z height to be 1-2 nm above a membrane plane. From these models, 18 starting Fab orientations were generated. A 70% POPC, 25% cholesterol, and 5% POPA coarse grain membrane was preassembled in a 30 ns simulation as describe in the pre-assembled pipeline. The assembled lipids and cholesterol molecules were extracted and placed into a 15x15x20 nm box along with a Fab molecule. This system was minimized in a vacuum using steepest descent algorithm, 30 fs time step, Berendsen thermostat at 300 K with time constant 1 $ps^{-1}$, and Berendsen barostat with isotropic coupling and time constant 12 $ps^{-1}$. Ions were added to 0.15 Mm final concentration and the system was then solvated with water molecules using a van der Waals radius of 0.21 nm. A minimization step was repeated for the solvated system. The 100 ns equilibration phase was performed with a 10 fs step. The velocity rescaling thermostat was used, with protein, lipid, and solvent independently coupled to a 300 K bath and a temperature coupling time constant of 2.0 $ps^{-1}$. The Berendsen barostat was used with semi-isotropic coupling and a pressure coupling time constant of 12 $ps^{-1}$. A 14 us production run was performed with 30 fs time step and velocity rescaling thermostat that coupled protein, lipid, and solvent independently to a 310 K bath with temperature coupling time constant of 1.0 $ps^{-1}$. The Berendsen barostat was used with semi-isotropic coupling and a pressure coupling time constant of 12 $ps^{-1}$.

**Headgroup Binding Site Characterization**

The x-ray structures of phosphate bound fab structures were used as references for alignments (PDB id: 4XCN, 6O3J, 5T80, 6SND for 4E10, PGZL1, 10E8, and LN01 respectively).

A metric to evaluate the accuracy to phosphate interaction in MD simulations and x-ray structures was calculated for every 0.5 ns of simulation time by superposing the interacting backbone residues of the MD simulation frame to interacting backbone residues from x-ray structure. For 4E10 and PGZL1, this superposition was done using CDRH1 residues 25-39. The RMSD was calculated between MD simulation's closest lipid headgroup phosphate coordinates and the phosphate in x-ray structure coordinates. For the 10E8 MD observed site, the FRL3 residues 83-87 (IMGT numbering) were aligned from every 0.5 ns of simulation time to the x-ray structure. After superposition, the root mean square distance (RMSD) was calculated between the closest lipid headgroup choline coordinates of the simulation frame and the phosphate coordinates in CDR-L1 loop of x-ray structure.

A bound state was then defined for each identified headgroup binding site. For 4E10 and PGZL1, a phosphate coordinate RMSD value and the distances of the interacting phosphate group to the loop backbone and side chain hydrogen atoms that were potential hydrogen bond donors were also measured. A phosphate group-hydrogen atom distance less than 5.25 Å was considered a potentially satisfied hydrogen bond interaction after accounting for thermal fluctuation and the mass averaged reference point for atoms of the phosphate group. For simulation frames where the phosphate coordinate RMSD was less than 2.0 Å (or 3.5 Å for the 10E8 MD predicted site) and at least two hydrogen bond interactions were considered potentially satisfied, that specific 0.5 ns interaction was deemed as a bound phosphate state.

For LN01 and LN01+MPER-TM x-ray site characterization, we aligned the MD frame CDR-L1 residues 23-35 to x-ray structures. A potential cation-pi cage interaction was defined as 2 or more distances between the closest lipid choline and center of aromatic rings for residues Tyr32, Tyr100G, Trp680, and Tyr681 being less than 5.5 Å. Polar interactions with a distance less than 5.25 Å between side chain hydroxyl groups of Thr30, Tyr32, Ser100F, and Tyr100G and closest lipid choline groups were also recorded. A hydrogen bond interaction was defined as a distance less than 5.25 Å between the Lys31 and the lipid phosphate. Headgroup occupancy was defined by satisfying one of the following conditions: a complete cation-pi cage formed around the choline headgroup, 2 or more satisfied interactions between lipid choline and nearby polar residues, or bivalent interaction including a hydrogen bond with lipid phosphate and one or more polar choline interaction or one or more cation-pi interactions. X-ray site RMSD was calculated with the MD lipid choline coordinates versus the x-ray structures lipid choline moiety.

The loading site interaction was also defined by aligning the MD frame CDR-L1 residues 23-35 to x-ray structure. A cation-pi cage was evaluated with the 2 or more distances less than 5.5 Å between lipid headgroup choline and residues Tyr49, Try52, Tyr100g, Trp100h and Tyr100i. Polar interactions between hydroxyl groups of Tyr49, Tyr52, Thr53, Thr100c, Ser100d, Tyr100g, and Tyr100i and lipid headgroup choline were defined with distances less than 5.25 Å. Lipid phosphate hydrogen bond interactions with Trp680, Lys683 and Ser100 were defined with a distances less than 5.25 Å. A loading site bound state was defined with similar logic to x-ray site occupancy and required satisfying one of the conditions listed above using the loading site specific residues described here. In LN01 systems, MPER-TM residues used in headgroup occupancy definitions were simply excluded from consideration. Loading site RMSD was calculated with the MD lipid choline coordinates versus a reference position of the averaged coordinates for bound choline headgroups.

**Antibody fragment RMSF calculations**

To calculate small scale changes in loop conformations, the MD simulation CDR-H1 residues were aligned from every 0.5 ns of MD simulated 4E10 and PGZL1 to relevant x-ray structures. FR-L3 residues from MD simulations to were aligned to 10E8 FRL3 x-ray structures, and CDR-L1 and CDR-H3 loops were aligned to LN01 x-ray structures by superposing the same residues described in headgroup RMSD calculations. A set of reference coordinates was calculated by averaging the loop or framework backbone atom coordinates across the aggregated of simulation time. We then calculated the RMSD of the MD frame backbone atoms to the average reference coordinates of backbone atoms after superposition. The RMSF values were subset into bound and unbound time based on headgroup occupancy across trajectories.

**Geometric Descriptor Calculations**

***Angles***

To calculate the angle to the membrane, two vectors along the pseudo-symmetry axes of the Fab and a normal vector to membrane plane were defined. The angle of approach vector is defined through two points: a point in the center of the tips of CDRL3 and CDRH3 loops, and a Fab center point between the proline residues (residue 40 with IMGT numbering) in heavy and light chains. The angle of rotation vector is defined with a point from the heavy chain hinge region through the light chain hinge region. The normal to the membrane plane is defined by fitting a plane to the phosphates in the top leaflet of the lipid bilayer and calculating a normal vector. Using these three vectors, the angle between the membrane plane and the antibody approach or rotation vector were calculated. All angles were transformed to be between -90 and 90 for angle of rotation and 0 and 90 for angle of approach.

***CDR Loop Depths***

The depth of CDR loops was calculated by finding the Z height of the mass averaged CDR loop center point (for L1, L2, L3, H1, H2, H3) and measuring the distance to the average Z height of the phosphates in the top lipid bilayer. CDR loop residues were selected based on IMGT numbering definitions.

***Membrane Embedded Surface Area***

The membrane embedded surface area was calculated by measuring a fully solvated Fab solvent accessible surface area (SASA) and subtracting the membrane embedded Fab SASA every 0.5 ns of atomistic simulation time. SASA was calculated with VMD measure plugin which implements a point based random sampling approach with 1.4 nm sphere.

***Interaction Profiles***

Interaction profiles were generated by defining the depth of each residue relative to the lipid bilayer over every 0.5 ns for atomistic simulations or every 30 ns for coarse grain simulations. The phosphate layer depth range was defined by calculating the average Z height of the top leaflet phosphates and expanding +/- 3 Å to account for headgroup size and thermal fluctuations in space. The glycerol layer was defined from the bottom of the phosphate layer – 4 Å based on glycerol molecule heights. The hydrocarbon layer was defined as up to 8 Å angstroms below the bottom of the glycerol layer. Because of the four to one mapping of atoms to beads in coarse grain representations, the layer definitions were expanded by 4 Å for coarse grain interaction profiling. Only the top leaflet layers were defined as a Fab was not observed to embedded deeper than below the top leaflet. Any non-lipid interacting residues were mapped to the solution layer of the system. The position of each residue in the Fab was mapped to a defined layer in the system based on residue Z height and time in each layer was aggregated over the total time of interest.

***Spontaneous Membrane Association***

To define an antibody that associated with the membrane in coarse grain systems, the distance of each Fab reside to the phosphate plane in the top or bottom bilayer was measures. This allowed us to account for a Fab that may diffuse across periodic boundaries in the defined system box. Tracking Fab residue positions every 30 ns, if a residue was within 4 Å of the phosphate layer, it was considered a Fab-membrane association.

**Structural Bioinformatics**

The PICES server was used to curate a list of 33,372 x-ray structure PDB entries with resolution below than 2.5 Å and less than 90% sequence similarity were downloaded into an in-house database. Method of Accelerated Search for Tertiary Ensemble Representatives (MASTER) was used to query protein structural fragments against the custom representative database to return hits with a maximum backbone RMSD of 1.8Å . These backbone matches were then re-queried to find hits containing sulfate or phosphate anions within 1.5 Å RMSD from the initial query phosphate.

**Clustering and medoid definitions**

Feature-based clustering was performed on the atomistic or coarse grain simulations. An 8-feature vector was defined for every 0.5 ns in atomistic simulation time that was composed of angle of approach, angle of rotation, and CDRHL1, CDRL2, CDRL3, CDRH1, CDRH2 and CDRH3 loop center of mass depths. In these clustering approaches, hierarchical agglomerative clustering with Ward's linkage method was used to minimize the variance between clusters defined. Euclidian distance thresholds were selected for each clustering experiment (antibody system and simulation scheme) to define clusters that optimize (macroscopic substates) or minimize (microscopic substates) divergence for average geometric descriptors across clusters in the context total clustering space.

To reduce the computational cost of clustering coarse grain simulation data, the feature vector was reduced to only include angle of rotation and angle of approach and sampled every 30 ns of simulation time. From here the same clustering methods as described before were applied and defined Euclidian distance cutoffs to generate 3 clusters from the sampled time for coarse grain systems., applied K-medoids clustering with an n equal to 1 was used to identify medoids that are the best representative snapshots of clusters.

**Coarse grain to all-atom backmapping**

Spontaneous insertion coarse grain clusters for 4E10, PGZL1, and 10E8 were defined using vectors describing each frame by angle of approach and angle of rotation. Hierarchical agglomerative clustering was performed with Ward's linkage method was used to minimize the variance between clusters. Euclidian distance thresholds were selected for so that three clusters were defined in for each antibody. Representative medoid frames were selected by calculating the frame that minimizes within-cluster sum of squares with Kmedoids algorithm from scikit-learn Clustering module.

To revert medoid frames from spontaneously associated coarse grain substates into atomistic representations, the original atomic Fab model (PDB ids: 2FX9, 6O3D, 5T85 for 4E10, PGZL1, and 10E8 respectively) was oriented relative to an assembled atomistic membrane (70% POPC, 25% Chol, 5% POPA) based on the corresponding coarse grain geometric representation by transforming the angle of approach and angle of rotation vectors. After docking the Fab, clashing lipid molecules were removed. Systems were solvated with TIP3 water molecules and neutralized with ions to a total concentrations of 0.15 mM NaCl. Minimization, equilibration, and production simulations were performed as previously described for atomistic simulations.

**Constant velocity pulling simulations**

To generate starting conformation ensembles, atomic Fab models were docked into HIV-like lipid bilayers with the Orientations of Proteins in Membranes (OPM) Positioning of Proteins in Membrane 2.0 (PPM) server. 500 ns equilibration simulations were performed as described in Atomistic Simulations and frames were extracted every 50 ns. The same Fab models were used for 4E10, PGZl1, and 10E8 as in primary atomistic simulations. A PGZL1 intermediate model was constructed by building loops into PDB ID 6O41 with ModLoop. PGZL1 germline model was built by introducing mutations into the PGZL1 intermediate model in PyMOL. Both PGZL1 germline and PGZL1 intermediate models were simulated for 50ns in solution with the same protocol in Atomistic Simulations prior to docking into bilayers and 500 ns equilibrations.

For pulling trajectories, starting frames were simulated with 1000 kJ/mol/nm$^3$ restraint on lipids for 100 ns production runs. The Berendsen thermostat was applied to protein, lipid, and solvent groups independently coupled to a 310 K bath with temperature time constant 1.0 ps$^{-1}$. The Berendsen barostat was used with semi-isotropic coupling and a 5.0 ps$^{-1}$ pressure time constant. Center of mass pulling was turned on between protein and lipid groups. Umbrella sampling potential was applied between the two groups at a rate of 0.03 nm/ns with a force constant of 1000 kJ/mol/nm in the direction away from the bilayer. Statistical comparisons for comparing max forces across systems were using with Tukey's multiple comparison test and unpaired t-tests.
